# The Impact of SIV-Induced Immunodeficiency on Clinical Manifestation, Immune Response, and Viral Dynamics in SARS-CoV-2 Coinfection

**DOI:** 10.1101/2023.11.15.567132

**Authors:** Alexandra Melton, Lori A Rowe, Toni Penney, Clara Krzykwa, Kelly Goff, Sarah Scheuermann, Hunter J Melton, Kelsey Williams, Nadia Golden, Kristyn Moore Green, Brandon Smith, Kasi Russell-Lodrigue, Jason P Dufour, Lara A Doyle-Meyers, Faith Schiro, Pyone P Aye, Jeffery D Lifson, Brandon J Beddingfield, Robert V Blair, Rudolf P Bohm, Jay K Kolls, Jay Rappaport, James A Hoxie, Nicholas J Maness

## Abstract

Persistent and uncontrolled SARS-CoV-2 replication in immunocompromised individuals has been observed and may be a contributing source of novel viral variants that continue to drive the pandemic. Importantly, the effects of immunodeficiency associated with chronic HIV infection on COVID-19 disease and viral persistence have not been directly addressed in a controlled setting. Here we conducted a pilot study wherein two pigtail macaques (PTM) chronically infected with SIVmac239 were exposed to SARS-CoV-2 and monitored for six weeks for clinical disease, viral replication, and viral evolution, and compared to our previously published cohort of SIV-naïve PTM infected with SARS-CoV-2. At the time of SARS-CoV-2 infection, one PTM had minimal to no detectable CD4+ T cells in gut, blood, or bronchoalveolar lavage (BAL), while the other PTM harbored a small population of CD4+ T cells in all compartments. Clinical signs were not observed in either PTM; however, the more immunocompromised PTM exhibited a progressive increase in pulmonary infiltrating monocytes throughout SARS-CoV-2 infection. Single-cell RNA sequencing (scRNAseq) of the infiltrating monocytes revealed a less activated/inert phenotype. Neither SIV-infected PTM mounted detectable anti-SARS-CoV-2 T cell responses in blood or BAL, nor anti-SARS-CoV-2 neutralizing antibodies. Interestingly, despite the diminished cellular and humoral immune responses, SARS-CoV-2 viral kinetics and evolution were indistinguishable from SIV-naïve PTM in all sampled mucosal sites (nasal, oral, and rectal), with clearance of virus by 3-4 weeks post infection. SIV-induced immunodeficiency significantly impacted immune responses to SARS-CoV-2 but did not alter disease progression, viral kinetics or evolution in the PTM model. SIV-induced immunodeficiency alone may not be sufficient to drive the emergence of novel viral variants.

## Introduction

The global outbreak of Coronavirus disease 2019 (COVID-19), caused by the highly infectious severe acute respiratory syndrome coronavirus 2 (SARS-CoV-2), has posed a significant and urgent public health challenge. First identified in Wuhan, China, in December 2019, the outbreak quickly spread to other countries across the globe. As of September 2023, the World Health Organization (WHO) has reported over 770 million global cases and nearly 7 million deaths [1]. While the majority of cases are asymptomatic or exhibit only mild symptoms, some individuals develop severe complications such as pneumonia, systemic inflammation, and coagulopathy, which can lead to organ failure, shock, and death [2–7]. Certain factors, such as a compromised immune system, advanced age, and comorbidities, such as cardiovascular disease, diabetes, and obesity, increase the risk of developing severe disease [8,9].

People living with HIV (PLWH) face an increased risk of several of these conditions, including a compromised immune system and a higher prevalence of cardiovascular disease. Additionally, PLWH have increased susceptibility to opportunistic infections such as pneumocystis pneumonia, which is the most common respiratory infection in patients with AIDS [10–12]. PLWH also experience elevated levels of inflammation, which significantly contributes to the development of severe respiratory disease, thromboembolisms, and other adverse outcomes associated with COVID-19 [13–15]. This raises concerns about the impact of HIV on the severity and persistence of SARS-CoV-2 infections. Studies examining whether HIV increases the risk of severe COVID-19 have yielded conflicting results. Initial studies indicated that PLWH had similar or even better outcomes [16–18] compared to those without HIV. However, larger population-based studies suggest that PLWH experience higher hospitalization rates and COVID-19-related deaths compared to the general population [19–23]. More recent research has suggested that unsuppressed viral loads or low CD4+ T cell counts are linked to suboptimal adaptive immune responses to SARS-CoV-2, affecting both T cell and humoral responses [24,25].

In addition to the concern of increased severity, HIV-associated immunodeficiency could potentially facilitate SARS-CoV-2 persistence and evolution, leading to the emergence of new variants of concern. A recent study by Karim et al. highlighted a case of an individual with advanced HIV who exhibited prolonged SARS-CoV-2 shedding with high viral loads and the emergence of multiple viral mutations [26]. While retrospective studies have explored the effects of HIV status on COVID-19 incidence and severity, controlled studies are lacking. To explore the feasibility of using an NHP model to address these gaps, we conducted a pilot study involving two pigtail macaques (PTM) chronically infected with SIVmac239. We exposed them to SARS-CoV-2 and monitored the animals for six weeks for clinical disease, viral replication, and viral evolution. Additionally, we performed detailed analyses of innate and adaptive immune responses, utilizing flow cytometry, cytokine/chemokine analysis, antibody binding and neuralization assays, and longitudinal single-cell RNA sequencing (scRNA-Seq) of bronchoalveolar lavage (BAL) cells following SARS-CoV-2 infection. We compared our findings with data from our previously published cohort of SIV-naïve, SARS-CoV-2-infected PTMs [27].

Despite the marked decrease in CD4+ T cells in the SIV+ animals prior to exposure to SARS-CoV-2, we found that disease progression, viral persistence, and evolution of SARS-CoV-2 were comparable to the control group. Overall, our findings suggest that SIV-induced immunodeficiency alters the immune response to SARS-CoV-2 infection, leading to impaired cellular and humoral immunity. However, this impairment does not significantly alter the course of infection. These findings contribute to a deeper understanding of the interplay between immunodeficiency and SARS-CoV-2 infection and propose a valuable model for evaluating vaccine and therapeutic strategies for immunocompromised individuals.

## Materials and Methods

### Research Animals

Two female pigtail macaques (PTM, Table 1) were inoculated intravenously with SIVmac239 (100 TCID50), followed by intranasal (0.5 mL per nare) and intratracheal (1 mL) administration of SARS-CoV-2 (1.1×10^6^ PFU/mL, USA WA1/2020) approximately one year later. Animals were monitored for six weeks following SARS-CoV-2 inoculation. Blood, bronchoalveolar lavage (BAL), and endoscopic gut biopsies were collected before and after SIVmac239 infection. Sampling pre- and post-SARS-CoV-2 infection included blood, BAL, and mucosal swabs (nasal, pharyngeal, and rectal). Physical examinations were performed throughout the course of the study. At the end of the study, complete postmortem examinations were performed with collection and histopathologic evaluation of 43 different tissues including all major organs and sections from each major lung lobe.

**Table 1.** Cohort of PTM used in this study.

| Animal ID | Sex | Age (y) | Weight (kg) at time of SIV inoculation | Weight (kg) at time of SARS-CoV-2 inoculation |
| --- | --- | --- | --- | --- |
| NV18 | Female | 3.82 | 4.45 | 4.45 |
| NV19 | Female | 4.67 | 5.76 | 5.85 |

### Ethics Statement

Pigtail macaques used in this study were purpose bred at Johns Hopkins University and moved to Tulane National Primate Research Center (TNPRC) for these experiments. Macaques were housed in compliance with the NRC Guide for the Care and Use of Laboratory Animals and the Animal Welfare Act. Animal experiments were approved by the Institutional Animal Care and Use Committee of Tulane University. The TNPRC is fully accredited by AAALAC International (Association for the Assessment and Accreditation of Laboratory Animal Care), Animal Welfare Assurance No. A3180-01. Animals were socially housed indoors in climate-controlled conditions with a 12/12-light/dark cycle. All the animals on this study were monitored twice daily to ensure their welfare. Any abnormalities, including those of appetite, stool, behavior, were recorded and reported to a veterinarian. The animals were fed commercially prepared monkey chow twice daily. Supplemental foods were provided in the form of fruit, vegetables, and foraging treats as part of the TNPRC environmental enrichment program. Water was available at all times through an automatic watering system. The TNPRC environmental enrichment program is reviewed and approved by the IACUC semi-annually.

Veterinarians at the TNPRC Division of Veterinary Medicine have established procedures to minimize pain and distress through several means. Monkeys were anesthetized with ketamine-HCl (10 mg/kg) or tiletamine/zolazepam (3-8 mg/kg) prior to all procedures. Preemptive and post procedural analgesia (buprenorphine 0.03 mg/kg IM or buprenorphine sustained-release 0.02 mg/kg SQ) was required for procedures that would likely cause more than momentary pain or distress in humans undergoing the same procedures. The animals were euthanized at the end of the study using methods consistent with recommendations of the American Veterinary Medical Association (AVMA) Panel on euthanasia and per the recommendations of the IACUC. Specifically, the animals were anesthetized with tiletamine/zolazepam (8 mg/kg IM) and given buprenorphine (0.01 mg/kg IM) followed by an overdose of pentobarbital sodium. Death was confirmed by absence of respiration, cessation of heartbeat, pupillary dilation, and lack of corneal reflex. The TNPRC policy for early euthanasia/humane endpoint was included in the protocol in case those circumstances arose.

### Isolation and Quantification of SIVmac239

Plasma SIVmac239 viral RNA (vRNA) extraction and quantification were performed essentially as previously described [28].

### Isolation of SARS-CoV-2 RNA

SARS-CoV-2 vRNA was isolated from BAL supernatant (200 µL) and mucosal swabs (nasal, pharyngeal, and rectal) using the Zymo Quick-RNA Viral Kit (Zymo Research, USA) as previously described [27,29]. Mucosal swabs, collected in 200 µL DNA/RNA Shield (Zymo Research, USA), were placed directly into the Zymo spin column for centrifugation to ensure complete elution of the entire volume. The Roche high pure viral RNA kit (Roche, Switzerland) was used to isolate vRNA from plasma (200 µL) per the manufacturer’s protocol. After isolation, samples were eluted in 50 µL DNase/RNase-free water (BAL and mucosal swabs) or Roche elution buffer (plasma) and stored at −80℃ until viral load quantification.

### Quantification of SARS-CoV-2 RNA

The quantification of SARS-CoV-2 RNA was performed according to methods previously described [27,29]. Genomic vRNA was quantified using CDC N1 primers/probe to determine the total amount of vRNA present. Additionally, primers/probe specific to nucleocapsid subgenomic (SGM) vRNA were utilized to estimate the quantity of replicating virus.

### Meso Scale Panels

To measure concentrations of various chemokine and cytokine protein targets, three V-plex MSD Multi-Spot Assay System kits were utilized: Chemokine Panel 1 (Eotaxin, MIP-1τ3, Eotaxin-3, TARC, IP-10, MIP-1α, IL-8, MCP-1, MDC, and MCP-4), Cytokine Panel 1 (GM-CSF, IL-1α, IL-5, IL-7, IL-12/IL-23p40, IL-15, IL-16, IL-17A, TNF-τ3, and VEGF-A), and Proinflammatory Panel 1 (IFN-γ, IL-1τ3, IL-2, IL-4, IL-6, IL-8, IL-10, IL-12p70, IL-13, and TNF-α) (Meso Scale Diagnostics, USA). Protein targets were measured in BAL supernatant (BAL SUP) and EDTA plasma following the manufacturer’s instructions, with an extended incubation time of overnight at 4°C to enhance sensitivity. Plasma samples were diluted 4-fold (Chemokine Panel 1) or 2-fold (Cytokine Panel 1 and Proinflammatory Panel 1) in the diluent provided in each kit. The plates were washed three times before adding prepared samples and calibrator standards. The plates were then sealed and incubated on a shaker at room temperature for two hours. Plates were immediately transferred to 4℃ for overnight storage. The following day, plates underwent three washes before the addition of detection antibody cocktails. Plates were then sealed and incubated on a shaker for two hours at room temperature. Following three final washes, MSD Read Buffer T was added to the plates, which were immediately read using a MESO QuickPlex SQ 120MM instrument (Meso Scale Diagnostics, USA). The concentration of each analyte was determined based on the standard curve plotted between known concentrations of calibrators and their respective signals. The Pheatmap package in R was used to generate the heatmap depicting log2 fold changes in chemokine and cytokine expression normalized to baseline (pre-SARS-CoV-2 inoculation).

### Isolation of Cells

SepMate-50 Isolation tubes (Stem Cell Technologies, Vancouver, Canada) were used according to the manufacturer’s protocol to isolate peripheral blood mononuclear cells (PBMCs) from whole blood. BAL samples were centrifuged at 1800 rpm at room temperature for 5 minutes. BAL supernatant was collected and stored at −80℃. BAL cell pellets were washed with PBS supplemented with 2% FBS. Tissue-specific lymphocytes were isolated from endoscopic duodenal pinches collected during the SIV portion of the study. Finely cut tissue pieces were added to a T-25 tissue culture flask and incubated in 25 mL Hanks Balanced Salt Solution (HBSS, Corning, USA) supplemented with 1mM EDTA (Invitrogen, USA) for 30 minutes at 37℃ at 400 rpm. After supernatant removal, samples underwent a second digestion in 25 mL RPMI (Gibco, USA) supplemented with 5% FBS, Collagenase II (60 units/mL, Sigma-Aldrich, USA), penicillin/streptomycin (100 IU/mL, Gibco, USA), 2 mM glutamine (Gibco, USA), and 25 mM HEPES buffer (Gibco, USA) for 30 minutes at 37℃ at 400 rpm. Samples were filtered through a 70-µm sterile cell strainer, washed, and resuspended in PBS with 2% FBS. Nexcelom’s Cellometer Auto 2000 (Nexcelom, USA) was used to count the cells. PBMCs were cryopreserved at approximately 1×10^7^ cells/mL in Bambanker cell freezing medium (GC Lymphotec, Japan).

### Flow Cytometry

Whole blood, thawed cryopreserved PBMCs, and freshly isolated cells from BAL and gut were washed with PBS supplemented with 2% FBS and stained with fluorescently labeled antibodies against markers listed in the Supplemental Section (S1 Table) as previously described [27]. Briefly, cells were incubated in Live/Dead stain cocktail (50 μL PBS + 0.5 μL live/dead stain per test) (Fixable Aqua Dead Cell Stain Kit, Invitrogen, Lithuania) in the dark for 20 minutes at room temperature. Cells were then washed and incubated in surface-stain cocktail containing 50 μL Brilliant Stain Buffer (BD Bioscience, USA) and antibodies listed in Supplemental Table 1. All samples were run on a BD FACSymphony A5 Cell Analyzer (BD Bioscience, USA), and data were analyzed with FlowJo 10.8.1 for Mac OS X (Tree Star, USA).

### T cell Cytokine Response to SARS-CoV-2

Mononuclear cells (MNCs) from blood and BAL were washed, pelleted, and resuspended in DMEM with 1% Anti-Anti and 10% FBS at 1×10^6^ cells/mL. Cells were stimulated overnight at 37℃, 5% CO_2_ with either cell stimulation cocktail (Biolegend, USA) or one of the following viral peptide pools obtained through BEI Resources, NIAID, NIH: Peptide Array, SARS Coronavirus Nucleocapsid Protein (NR-52419), Spike Glycoprotein (NR-52402), or Membrane Protein (NR-53822), along with co-stimulatory antibodies (CD28 and CD49d at 1 μL/mL) and Brefeldin-A (1 μL/mL, BioLegend, USA). LIVE/DEAD and surface staining was performed as described above. To measure cellular response to viral antigen, cells were washed in PBS containing 2% FBS, fixed and permeabilized with Cytofix/Cytoperm Buffer (BD Biosciences, USA). Cells were incubated in intracellular stain cocktail for 30 minutes at room temperature (S1 Table), washed with 1x BD Perm/Wash Buffer and fixed in 1x BD Stabilizing Fixative (BD Biosciences, Franklin Lakes, NJ).

Overnight stimulation, surface and intracellular staining of BAL cells isolated from SARS-CoV-2 infected animals were performed under BSL-3 safety conditions. Cells were fixed with 2% Paraformaldehyde for 60 minutes before removal from BSL-3. Samples were run on the BD FACSymphony and analyzed via FlowJo as described above.

### Meso Scale COVID-19 IgA and IgG Panels

V-PLEX COVID-19 serological assays were used to quantify serum levels of IgA and IgG binding antibodies to SARS-CoV-2 Spike, Spike N-Terminal Domain (S1 NTD), and Spike Receptor Binding Domain (S1 RBD) (Panel 1, Meso Scale Discovery, USA), following the manufacturer’s protocol. Briefly, plates were first incubated at room temperature on a shaker in MSD Blocking solution for 30 minutes, followed by 3 washes with 1X MSD Wash buffer. Plasma samples were diluted 100-(IgA kit) or 1000-fold (IgG kit) and plated in duplicate, along with controls and standards used to generate a seven-point calibration curve. Plates were then sealed and incubated at room temperature on a shaker for 2 hours. Following this, the plates were washed three times before addition of 1X detection antibody to each well. After a 1-hour incubation, plates were washed a final 3 times, and MSD GOLD Read Buffer B was added to the plates. Plates were read immediately using a MESO QuickPlex SQ 120MM instrument. The concentration of IgA and IgG antibodies was determined using the standard curve generated by plotting the known concentrations of the standards and their corresponding signals.

### SARS-CoV-2 Microneutralization (PRMNT) Assay

A microneutralization assay (PRMNT) adapted from Amanat et al. 2020 [30] was used to assess the presence of neutralizing antibodies in serum of SIV+ and SIV naïve SARS-CoV-2 infected PTMs. Vero/TMPRSS2 cells (JCRB Cell Bank, Japan) were seeded in 96-well tissue culture-treated plates to be subconfluent at the time of assay. Serum samples were diluted in dilution buffer (DMEM, 2% FBS, and 1% Anti-Anti) to an initial dilution of 1:5, followed by six 3-fold serial dilutions. SARS-CoV-2 (WA1/2020, BEI, USA) was diluted 1:3000 in dilution buffer and added in equal proportions to the diluted sera under Biosafety Level 3 (BSL-3) conditions. Samples were then incubated at room temperature for 1 hour. The culture media was removed from the 96-well Vero cell culture plates, and 100 µL of the virus/sera mixture was added to each well. Dilution buffer and diluted virus (1:6000) were used as the negative and positive controls, respectively. Plates were then incubated for 48 hours at 37°C and 5% CO_2_. After the incubation period, the medium was removed, and 100 µL of 10% formalin was carefully added to each well. The plates were allowed to fix overnight at 4°C before being removed from the BSL-3 facility.

The staining of the plates was conducted under BSL-2 conditions. After carefully removing the formalin, the cells were washed with 200 µL PBS, followed by the addition of 150 µL of permeabilization solution (0.1% Triton/PBS). Plates were then incubated at room temperature for 15 minutes. Following the incubation, the cells were washed with PBS and blocked with 100 µL of blocking solution (2.5% BSA/PBS) for 1 hour at room temperature. After removing the blocking solution, 50 µL of the primary antibody (SARS-CoV-2 Nucleocapsid Antibody, Mouse Mab, Sino Biologicals, #40143-MM08) diluted 1:1000 in 1.25% BSA/PBS was added to each well, followed by a 1-hour incubation at room temperature. The plates were then washed twice with PBS, decanted, and gently tapped on a paper towel to ensure complete antibody removal. Next, 100 µL of the secondary antibody, Goat anti-Mouse IgG (H+L) Cross-Adsorbed Secondary Antibody (Invitrogen, #A16072) diluted 1:3000 in 1.25% BSA/PBS was added to each well. The plates were incubated for 1 hour at room temperature. Following the incubation period, cells were washed as described above. To initiate color development, 100 µL of 1-Step Ultra TMB-ELISA developing solution (Thermo Scientific, #34028) was added to each well. The plates were then incubated in the dark at room temperature for 10 minutes. To stop the reaction, 50 µL of 1N sulfuric acid was added to each well. The optical density was measured and recorded at 450 nm on a Tecan Sunrise Microplate Reader (Tecan, Switzerland). The averages of the positive control wells and negative control wells were calculated separately, and percent inhibition was calculated for each well.

### Single-Cell RNA Sequencing (scRNAseq) of BAL Cells

For single-cell sequencing of bronchoalveolar lavage (BAL) cells, we collected samples before SARS-CoV-2 inoculation and on days 2, 7, 21, and 28 post-challenge. BAL samples were centrifuged at room temperature for 5 minutes at 1800 rpm, and the resulting cell pellets were resuspended in DMEM supplemented with 10% FBS and 1% Anti-Anti. We used the Parse Biosciences cell fixation kit following the manufacturer’s instructions for PBMCs to fix the cells (Parse Biosciences, USA). Specifically, we fixed 1 million cells per animal/timepoint in a 15 mL falcon tube. The fixed cells were stored at −20℃ until all samples were collected.

To enable multiplexing of samples, the Parse Single-cell whole transcriptome kit, which utilizes a combinatorial barcoding approach (Evercode WT, Parse Biosciences, USA), was employed. This allowed us to barcode and multiplex 10 samples collected from the coinfected animals across five timepoints. For analysis of the processed cells, we conducted two separate runs: the first run included approximately 15,000 cells, while the second run consisted of approximately 42,000 cells. The sublibraries from each run were pooled and sequenced on an Illumina NextSeq 2000 platform, yielding an average depth of 27,165 reads per cell for the first batch and 29,088 reads per cell for the second.

### Analysis of Single-Cell RNA Sequencing Data

For analysis of the single-cell sequencing data, we utilized the Parse Biosciences pipeline (v1.0.4.) to generate cell-gene matrix files using concatenated GTF annotations for the Rhesus macaque genome (Macaca mulatta, GCA_003339765.3), SARS-CoV-2 genome (GCA_009858895.3) and SIV genome (GenBank Accession # M33262.1).

Subsequently, the scRNAseq data analysis was performed using the Seurat package in R [31]. The *cell-gene matrix* (DGE.mtx), *cell metadata* (cell_metadata.csv), and *all genes files* (all_genes.csv) generated from both experimental runs using the Parse Biosciences pipeline were imported into R using the readMM and read.delim functions. Seurat objects were then created for each run, and the raw count matrices were merged using the merge command.

To ensure data quality, cells with more than 5% mitochondrial genes, fewer than 200 genes, or more than 2500 genes were excluded from further analysis. The data were normalized and scaled using the NormalizeData and ScaleData functions following the standard Seurat workflow. To account for batch effects and biological variability, we applied the Harmony [32] algorithm, which integrates the data by clustering cells based on their cell type rather than specific dataset conditions. Uniform manifold approximation and projection (UMAP) dimensional reduction was performed on the integrated Seurat object, using 20 dimensions based on the Harmony embeddings. Louvain clustering with a resolution of 0.5 was then conducted using the FindNeighbors and FindClusters functions to identify distinct cell clusters. After determining which cells contained SARS-CoV-2 or SIV transcripts, we excluded Day 2 samples from further analysis due to sample quality for one of the coinfected animals. After removing Day 2, we followed the same method as described above for quality control and integration.

### Identification of Cell Types

Cell type annotation was performed by identifying differentially expressed genes (DEGs) using the FindAllMarkers function, which utilizes the Wilcoxon rank-sum test, to determine significant differences in gene expression. Cell clusters were annotated based on expression of canonical cell marker genes. We identified 24 cell types, including epithelial cells (*TPPP3*), monocytes/macrophages (*MRC1, MARCO*), proliferating macrophages (*MRC1, MARCO, MKI67, HMGB2*), T cells (*CD3E*), proliferating T cells (*CD3E, MKI67, HMGB2),* invariant natural killer T cells (iNKT*, CD3E, IL7R*^hi^), natural killer cells (NK, *NKG2D*), Neutrophils (lineage negative), B cells (*MS4A1, CD19, CD79A*), plasma cells (*JCHAIN*), mast cells (*HPGE, CPA3, KIT*), plasmacytoid dendritic cells (pDC, *IRF8*), and myeloid dendritic cells (mDC, *ITGAX*). We used the subset function for subclustering analysis of monocyte/macrophage and T cell clusters. Again, the standard Seurat workflow for quality control and the Harmony algorithm for integration were applied. For the T cell subcluster, the number of dimensions was reduced to 10 in the RunUMAP function, and the resolution for FindClusters function was set to 0.2 to refine the clustering results.

### Differential Gene Expression and Gene Set Enrichment Analysis (GSEA) of Monocyte/Macrophage Subclusters

Differential gene expression analysis was conducted among the six monocyte/macrophage subclusters using the FindMarkers function in Seurat. Volcano plots were generated to visualize the results, highlighting genes with an average log2 fold change (log2fc) greater than 0.25 or less than −0.25 and a p-value less than 0.05 indicating statistical significance. GSEA was performed by ranking the list of DEGs based on their average log2fc. This ranking strategy enables the identification of pathways that show enrichment in our gene set, even when individual genes may not reach statistical significance. By considering the collective contribution of genes, we can uncover upregulated pathways that play a significant role in our analysis. Gene symbols were converted into Entrez IDs using the Metascape [33] website (https://metascape.org). We performed GSEA using the Hallmark [34] gene set from The Broad Institute Molecular Signature Database [35,36] (MSigDB). The msigdbr function was used to import the Hallmark gene set, and GSEA analysis was performed using the fgsea function [37]. Bar graphs were generated to illustrate the net enrichment scores (NES) of significantly enriched pathways within each subcluster using a false discovery rate (FDR) threshold of less than 0.1. The same strategy was applied for Hallmark GSEA, comparing NV18 and NV19 at baseline and 7-dpi with an increased FDR of 0.2. Additionally, we compared days 7, 21, and 28 to baseline for each animal and included both KEGG [38–40] and Hallmark gene sets for GSEA.

## Results

### Experimental Design and Viral Dynamics in SIV-infected Pigtail Macaques Prior to SARS-CoV-2 Exposure

Two female pigtail macaques (PTM, NV18 & NV19) were infected intravenously (iv) with SIVmac239 (100 TCID_50_) and monitored for approximately one year prior to exposure to SARS-CoV-2 (Wa1/2020, 2.2×10^6^ PFU, in/it) (Fig 1A). SIV viral dynamics in plasma followed the typical pattern, with peak viremia occurring approximately two weeks after infection, followed by a set point of around 1×10^6^ for NV18 and 1×10^5^ for NV19 (Fig 1B). The uncontrolled viremia led to a substantial progressive decrease in CD4+ T cells in all sampled compartments (plasma, BAL, and gut) (Fig 1C-E). Notably, beginning at approximately eight weeks post-SIV infection, NV18 exhibited few to no detectable CD4+ T cells in BAL and gut, and these levels remained persistently low until the time of SARS-CoV-2 exposure. The other animal, NV19 also experienced a decline in CD4+ T cells across all sampled compartments, and although levels began to rebound, they never returned to pre-infection levels.

**Figure 1.**
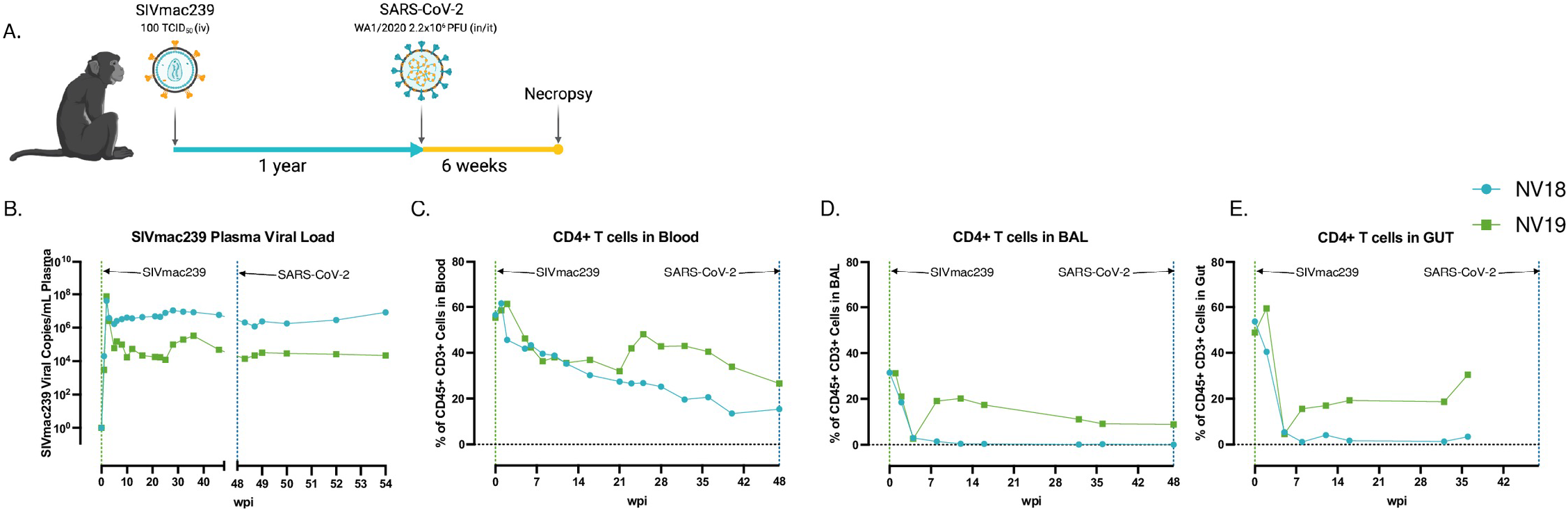
Before SARS-CoV-2 exposure, PTM experienced uncontrolled SIV viremia and immunodeficiency due to loss of CD4+ T cells. **A.** Overall study design. Two female pigtail macaques (PTM, NV18 & NV19) were inoculated with SIVmac239 (100 TCID_50,_ iv), followed by SARS-CoV-2 (Wa1/2020, 2×10^6^, in/it) challenge approximately one year later. Figure created with BioRender (https://BioRender.com). **B.** Quantification of SIVmac239 RNA levels in plasma overtime (Quantitative RT PCR). **C-E.** CD4+ T cell kinetics following SIVmac239 infection in blood (**C**), bronchoalveolar lavage (BAL) (**D**), and gut (**E**).

### Impact of SIV-Induced Immunodeficiency on SARS-CoV-2 Replication and Evolution

We then sought to investigate how SIV-induced immunodeficiency affects SARS-CoV-2 viral replication and evolution in our PTM model. We hypothesized that the observed immunodeficiency in the SIV-infected PTMs would enhance SARS-CoV-2 viral persistence, thereby increasing the risk of viral evolution. Using qRT-PCR, we tracked viral genomic (Fig 2A-E) and SGM (Fig 2F-J) RNA in mucosal swabs (nasal, pharyngeal, and rectal), BAL supernatant (sup), and plasma for six weeks. We compared viral dynamics in our coinfected animals with our previously published cohort of SIV-naïve PTMs [27]. Viral dynamics in BAL showed robust viral replication during acute infection in both the SIV+ and the controls with viral levels becoming undetectable in all animals by 21 days post infection (dpi). The coinfected animals cleared vRNA in the rectal mucosa by 14-dpi, the pharynx by 21-dpi, and the nasal mucosa by 28-dpi. The SIV-naïve animals had low levels of detectable virus in the nasal and rectal mucosa at their study end point of 21-dpi, with no detectable virus in the pharynx or plasma.

**Figure 2.**
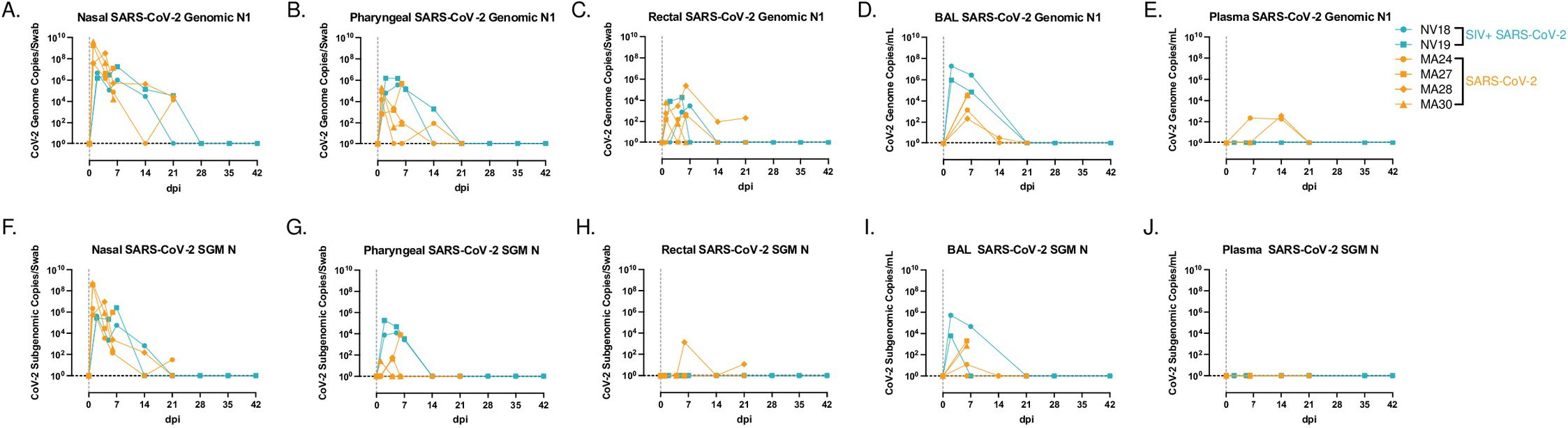
SARS-CoV-2 viral dynamics. NV18 and NV19 were inoculated with SARS-CoV-2 (1×10^6^ TCID50) 48 weeks post SIVmac239 infection through a combination of intranasal (in) and intratracheal (it) exposure, indicated as Day 0. **A-J.** Comparison of genomic (**A-E**) and subgenomic (SGM, **F-J**) SARS-CoV-2 mRNA levels in mucosal swabs (**A-C, F-H**), BAL supernatant (**D, I**) and plasma (**E, J**) in coinfected animals (blue) and a previously published cohort of SIV naïve PTM (orange).

Furthermore, we were unable to detect genomic or SGM vRNA in plasma in either of the coinfected animals. Surprisingly, both SIV+ animals cleared SARS-CoV-2, similar to the controls, and the absence of prolonged viral persistence consequently precluded any significant viral evolution, with H655Y being the only spike mutation detected in multiple samples from both coinfected animals at more than 25% of sequence read, including NV18 nasal and pharyngeal from day 2 and pharyngeal from day 5 and NV19 rectal sample from day 2. However, this mutation was also present at a low frequency in the inoculum, precluding any analysis of intrahost selection.

### Clinical Manifestations and Postmortem Observations in Coinfected PTM

Animals coinfected with SIVmac239 and SARS-CoV-2 were closely monitored for six weeks following SARS-CoV-2 inoculation. In line with clinical findings in our previous pigtail study, the coinfected animals exhibited only mild COVID-19 symptoms. This outcome was unexpected given that previous studies have indicated PLWH face a higher risk of severe disease attributed to factors such as low CD4+ T cell counts and uncontrolled viremia, both of which were observed in our SIV+ animals [19–23]. Similar to the controls, no significant changes in body weight, temperature, or blood oxygen saturation levels were observed in the coinfected animals (S1 Fig). Furthermore, thoracic radiographs of the coinfected animals closely resembled those of the control group, revealing only subtle changes consistent with mild to moderate manifestations of COVID-19 (S2 Fig). Upon postmortem examination, both animals demonstrated histopathologic changes consistent with chronic SIV infection. Neither animal had lesions that were attributed to SARS-CoV-2 infection, indicating that lesions had resolved. This resolution of SARS-CoV-2-associated lesions is expected given the six-week post-infection time point, viral clearance in these animals, and what has previously been reported in the NHP model. One animal, NV18, had an opportunistic Pneumocystis infection and SIV syncytial giant cells compatible with simian AIDS (SAIDS).

### Assessment of Cytokine and Chemokine Levels in Blood and BAL Following SARS-CoV-2 Infection of SIV+ PTM

To assess the changes in cytokine and chemokine levels in blood and BAL following SARS-CoV-2 infection in the coinfected animals, we utilized the MesoScale V-plex MSD Multi-Spot Assay System (Figs 3 and S3). Similar to findings by Huang et al. [41], who reported elevated plasma concentrations of MIP-1⍺, MCP-1, IL-7, IL-10, IP-10, IL-2, and GM-CSF, in hospitalized patients, we observed increased levels of MIP-1⍺, MCP-1, IL-7, and IL-10 in BAL supernatant from both coinfected PTMs at 2-dpi (Figs 3 and S3A-S3C). Additionally, at 7-dpi, the more immunocompromised animal (NV18) exhibited higher levels of IP-10, IL-2, and GM-CSF in BAL supernatant. We also detected increased plasma levels of MIP-1⍺ and MCP-1 for both animals at 2-dpi. Notably, pulmonary levels of several cytokines and chemokines exhibited a secondary increase at days 21 or 28 post-infection (MIP-1⍺, MCP-1, IL-7, IL-10, IP-10, IL-2, and GM-CSF). Overall, NV18 exhibited higher levels of these markers in the lung, while NV19 tended to have higher levels in the plasma.

**Figure 3.**
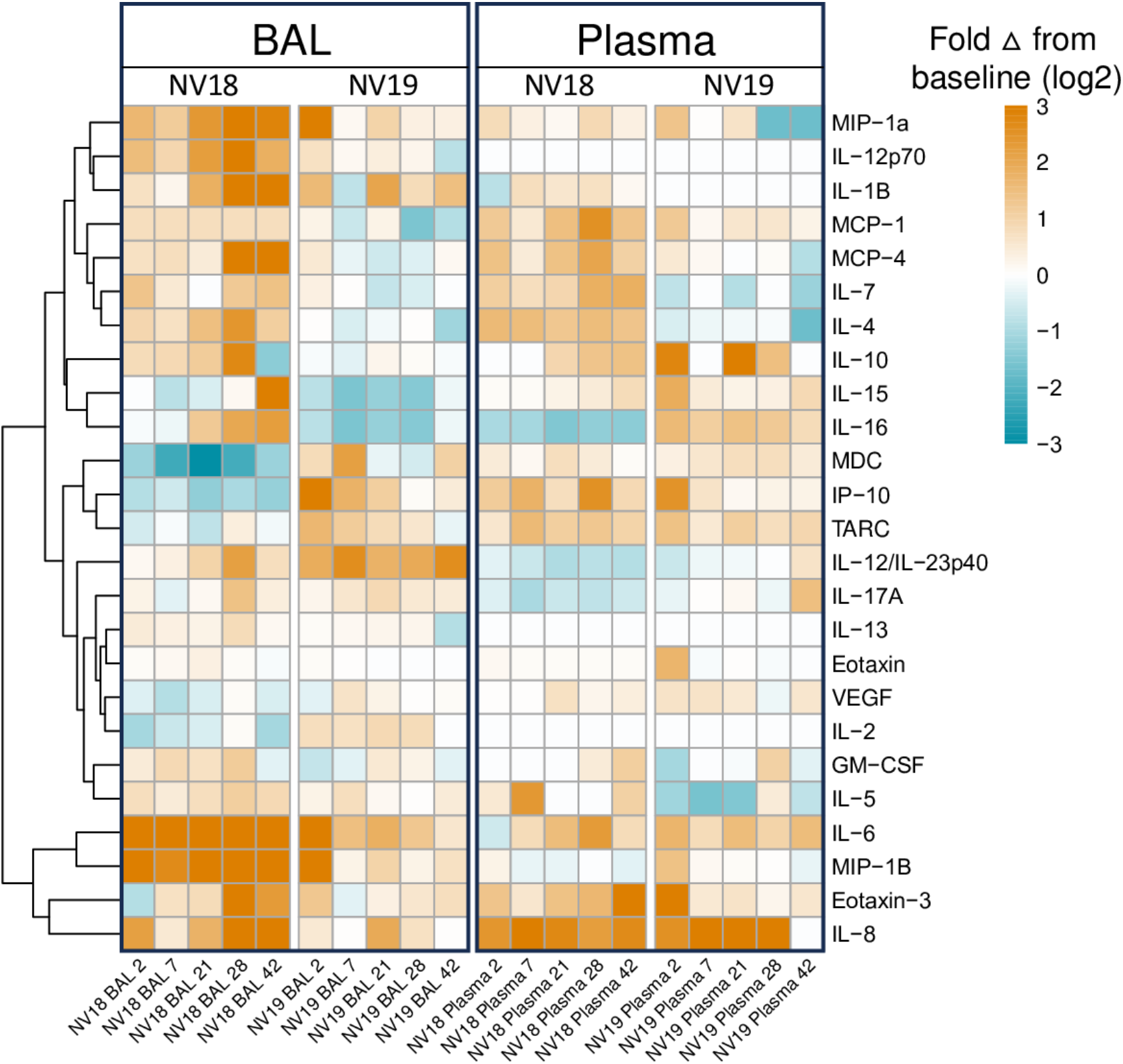
Meso Scale analysis of cytokine and chemokine fluctuations in blood and BAL in PTM coinfected with SIV and SARS-CoV-2. **A.** Heatmap indicating changes in cytokine and chemokine levels in BAL supernatant and plasma. Data represents log2 fold change from baseline (pre-SARS-CoV-2 infection). **B-YY.** Line graphs illustrating cytokine, proinflammatory cytokine, and chemokine dynamics in BAL supernatant (**B-H, P-W, FF-YY**) and plasma (**I-O, X-EE, PP-YY**) before and 2-, 7-, 21-, 28-, and 42-days post SARS-CoV-2 infection.

### Pulmonary Monocyte Infiltration and Chemokine Dynamics in SARS-CoV-2 Infection

Pulmonary inflammatory immune cell infiltration, particularly by monocytes/macrophages, is well characterized in COVID-19 [27,29,42–44]. Chemokines associated with monocyte recruitment in the blood include IP-10, TARC/CCL17, and MCP-1/CCL2 [43]. In our study, we observed elevated levels of these chemokines in plasma at 2-dpi in both animals, accompanied by an increase in monocytes (Figs S3C and 4). Furthermore, NV19 demonstrated increases in these chemokines in BAL supernatant at 2-dpi. Interestingly, the more immunocompromised animal (NV18) initially exhibited a decrease in IP-10 and TARC levels in the lung, followed by a rise at day 7.

We also observed pulmonary infiltration of classical monocytes at 2-dpi (NV18 and NV19, Fig 4B) and intermediate monocytes at days 2 (NV18) and 7 (NV19) (Fig 4C). Similar monocyte kinetics were observed in the SIV-naïve animals, with a transient increase in pulmonary infiltrating monocytes during the acute phase of SARS-CoV-2 infection, returning close to baseline at approximately 14-dpi. While monocyte kinetics were similar for NV19 and the control animals, NV18 had higher peripheral levels of all monocyte subsets, along with increased pulmonary monocytes at 21 and 28-dpi.

**Figure 4.**
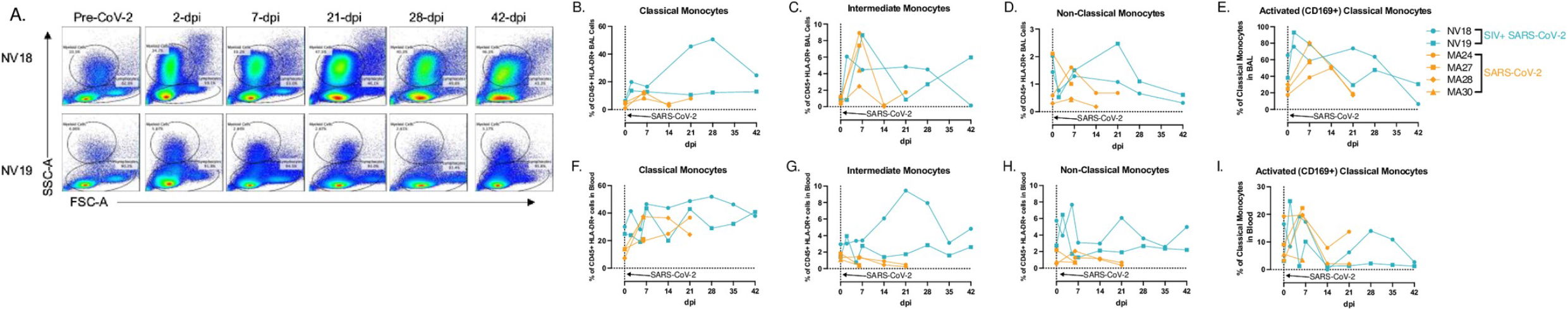
Monocyte/macrophage kinetics in BAL and blood following SARS-CoV-2 infection of SIV+ and SIV naïve PTM. **A.** Representative flow cytometry dot plots of pulmonary infiltrating myeloid cells and lymphocytes in the lungs of two SIV+ PTM before and 2, 7, 21, 28, and 42 days after SARS-CoV-2 exposure. Gated on Time>Live>Single cells>CD45+>SSC-A vs FSC-A. **B-E, F-I.** Frequencies of Classical (CD45+ HLA-DR+ CD14+ CD16-) (**B,F**) intermediate (CD45+ HLA-DR+ CD14+ CD16+) (**C,G**), and non-classical monocytes (CD45+ HLA-DR+ CD14-CD16+) (**D,H**) in BAL (**B-D**) and blood (**F-H**) before and after SARS-CoV-2 infection. Day 0 = day of SARS-CoV-2 infection. **E,I.** Activated Classical Monocytes (CD169+) in BAL and blood. Figures B-I: SIVmac239/SARS-CoV-2 infected PTM (blue), and SARS-CoV-2 only infected PTM (orange).

Notably, we also observed a spike in several cytokines (IL-6, 1L-10, IL-13, 1L-2, 1L-4, IL-12p70, IL-1B, IL-16, IL-17A, VEG-F, GM-CSF, IL-5, and IL-7) and chemokines (TARC, IL-8, MIP-1B, Eotaxin, Eotaxin-3, MCP-4, MIP-1a, and MIP-1a) in BAL at 28-dpi, suggesting a potential role for these markers in monocyte recruitment and/or function (S3 Fig). It is important to note, however, that due to sample availability constraints, we were unable to conduct the same Meso Scale cytokine/chemokine analyses on the SIV-naïve animals. Given that we lack direct comparison to the SIV-naïve cohort, we are limited in our ability to draw definitive conclusions from our cytokine and chemokine results.

### T cell Dynamics in Blood and BAL Following SARS-CoV-2 Infection

T lymphopenia, specifically of CD4+ T cells, is a common feature observed in human COVID-19 patients. This, compounded with low CD4+ T cell counts due to advanced HIV/SIV infection, may delay the clearance of SARS-CoV-2, increase the risk of viral evolution, and promote disease progression [45,46]. In our study, both coinfected animals displayed signs of immunodeficiency with a substantial loss of CD4+ T cells in blood, lung, and gut prior to SARS-CoV-2 exposure (Fig 1C-E). Acutely following SARS-CoV-2 infection, both animals experienced a further decline in peripheral CD4+ T cells. In NV19, this decline was transient and reached a nadir at 2-dpi. However, in the more immunocompromised animal, NV18, the loss persisted, and CD4+ T cells remained undetectable in both blood and BAL for the remainder of the study (Figs 5A, 5C, 5E, and 5G). Both animals showed a reduction in the overall CD3+ T cell population in BAL at 2-dpi with levels returning to baseline in NV19 at 7-dpi (Fig 5F). T cell dynamics in the SIV-naïve animals exhibited patterns similar to those of NV19, though with slightly delayed kinetics (Figs 5B-D and 5F-H). Despite the loss of CD4+ T cells, both coinfected animals successfully cleared SARS-CoV-2, suggesting the involvement of innate immune mechanisms in controlling the infection.

**Figure 5.**
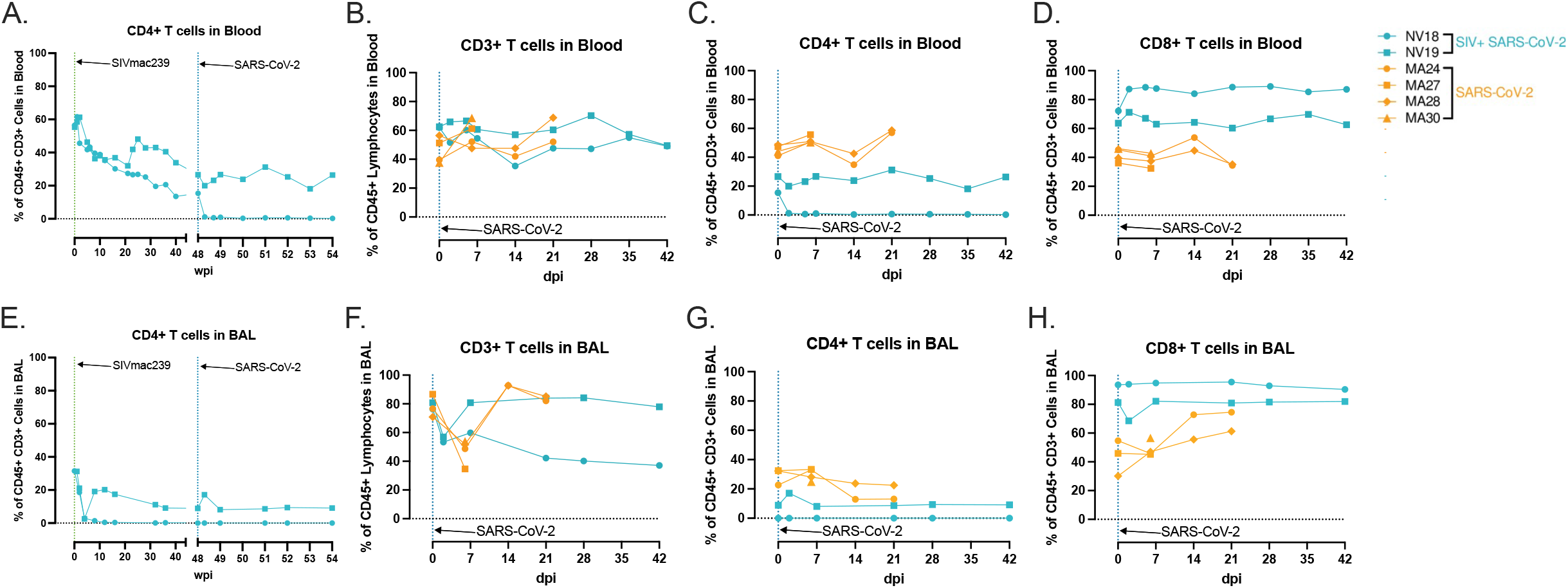
T cell dynamics in blood and BAL following SARS-CoV-2 infection of SIV+ and SIV naïve PTM. CD4+ T cell kinetics in blood **(A)** and BAL **(G**), following SIVmac239 and SARS-CoV-2 infection. **H-L.** Peripheral and pulmonary T cell dynamics in SIV+ SARS-CoV-2 coinfected PTM (blue). Historical data from four SIV naive SARS-CoV-2 infected PTM in orange. Comparison of overall CD3+ T cell populations (**B&F**) as well as T cell subsets; CD4+ (**C&G**), CD8+ (**D&H**).

### Diminished Cellular Immune Response to SARS-CoV-2 in Coinfected Animals with Severe T Cell Lymphopenia

To evaluate the cellular immune response to SARS-CoV-2 infection, we stimulated mononuclear cells isolated from BAL with peptides derived from SARS-CoV-2 Spike, Membrane, or Nucleocapsid proteins and assessed cytokine responses using flow cytometry. In our previous PTM study, we showed that at 21-dpi, the SIV-naïve animals developed pulmonary CD4+ and CD8+ SARS-CoV-2-specific T cell responses, that were predominately CD4 driven. However, in our current study, neither coinfected animal had detectable virus-specific cellular immune responses to peptide stimulation (Fig 6). Consistent with our previous findings, we were unable to detect virus-specific T-cell responses in the blood at 21-dpi (S4 Fig). Our findings show that severe CD4+ T-cell lymphopenia, resulting from advanced SIV infection, significantly impairs the cellular immune response to SARS-CoV-2 in the lungs.

**Figure 6.**
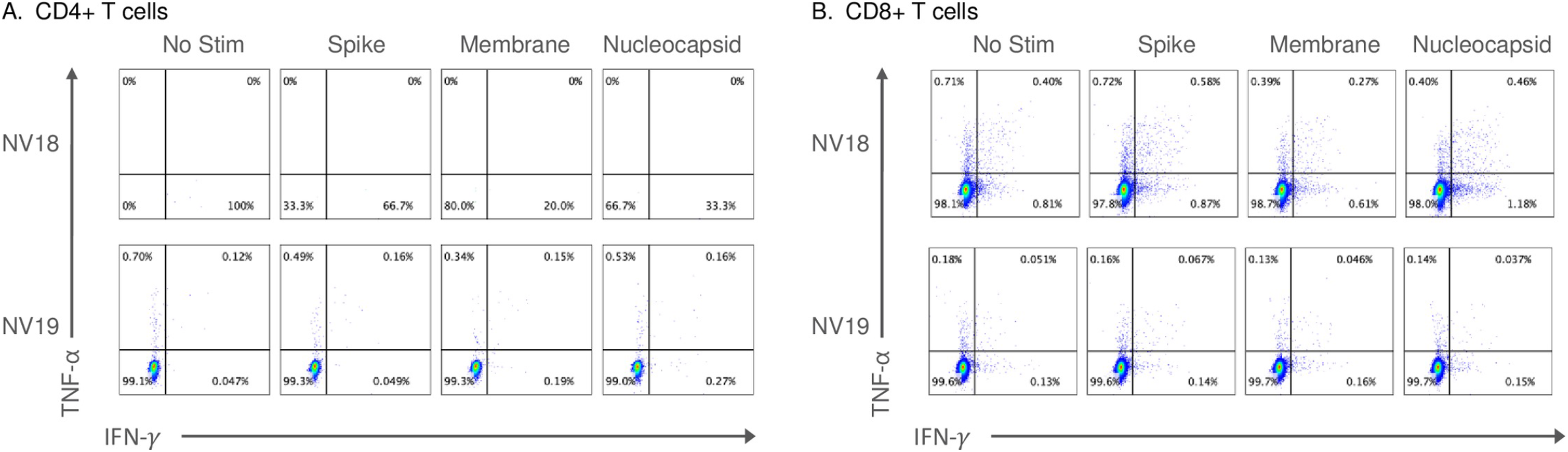
SARS-CoV-2 specific T cell responses were undetectable in the lung 21-days-post-infection. Two female pigtail macaques (PTM, NV18 & NV19) co-infected with SIVmac239 and SARS-CoV-2 shown. **A&B**. Flow cytometry dot plots demonstrating the IFN-γ and TNF-⍺ response of CD4+ (**A**) and CD8+ (**B**) T cells to overnight SARS-CoV-2 peptide (spike, membrane, and nucleocapsid) stimulation. No Stim = cells incubated overnight without peptide stimulation.

### Impaired Humoral Immune Response to SARS-CoV-2 Infection

We then aimed to assess neutralization capacity of serum antibodies using a microneutralization assay (PRMNT) [30]. By 14-dpi, the SIV-naïve animals demonstrated detectable neutralizing antibodies against SARS-CoV-2, whereas the coinfected animals failed to generate a neutralizing antibody response (Fig 7A). Additionally, using the V-PLEX COVID-19 serological assay kit from Meso Scale Discovery, we measured IgA and IgG binding antibody levels in serum. By 21-dpi, we detected IgA (Fig 7B) and IgG (Fig 7C) binding antibodies targeting various domains of the Spike protein, including the receptor binding domain (RBD), Spike S1 and S2 domains, and the Spike N-terminal domain (NTD) in the SIV-naïve PTMs. However, we were unable to detect IgA or IgG binding antibodies in the serum of the coinfected animals. Our data demonstrate that the coinfected animals failed to generate virus-specific T cell and humoral immune responses highlighting the impact of pre-existing immunodeficiency on the development of adaptive immunity during coinfection.

**Figure 7.**
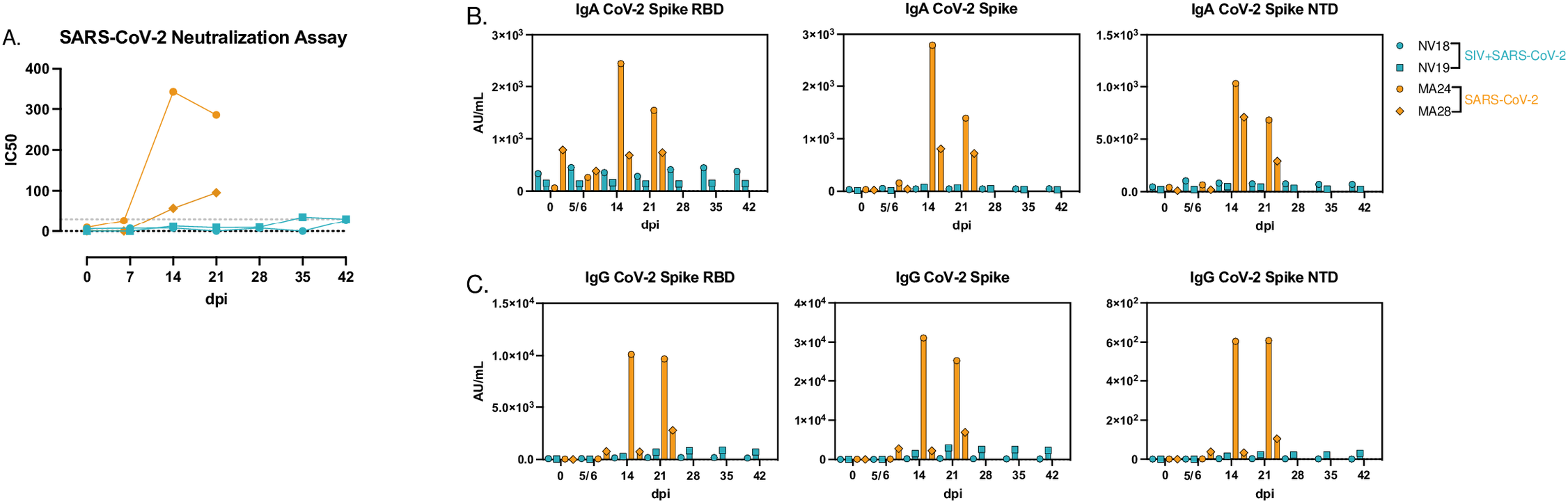
Humoral immune response to SARS-CoV-2 infection. **A.** SARS-CoV-2 neutralization assay depicting serum antibody levels against SARS-CoV-2 using Vero TMPRSS2 cells. **B&C.** MesoScale analysis of IgA (**B**) and IgG (**C**) binding antibodies to SARS-CoV-2 Spike receptor binding domain (RBD), Spike glycoprotein 1 and 2 (S1&S2), and Spike N-Terminal Domain (NTD).

### Single-Cell RNA Sequencing

To gain a more detailed understanding of the pulmonary immune response to SARS-CoV-2 infection in the coinfected animals, we conducted single-cell RNA sequencing (scRNAseq) on cells isolated from BAL at baseline (48 weeks post SIVmac239 exposure) and days 2, 7, 21, and 28 post-SARS-CoV-2 infection (Fig 8). This approach allowed us to examine the immune response at a higher resolution and capture dynamic changes over the course of coinfection (Figs 8C and 8D). Cell type clusters identified in BAL included monocytes/macrophages, dendritic cells (DC), neutrophils, natural killer (NK) cells, invariant natural killer T (iNKT) cells, T cells, B cells, plasma cells, mast cells (MC), as well as proliferating T cells and macrophages (Figs 8A, 8B, and S5). It should be noted that due to sample availability, we were unable to perform scRNAseq analysis on the SIV-naïve animals therefore the results presented here within should be interpreted as observational.

**Figure 8.**
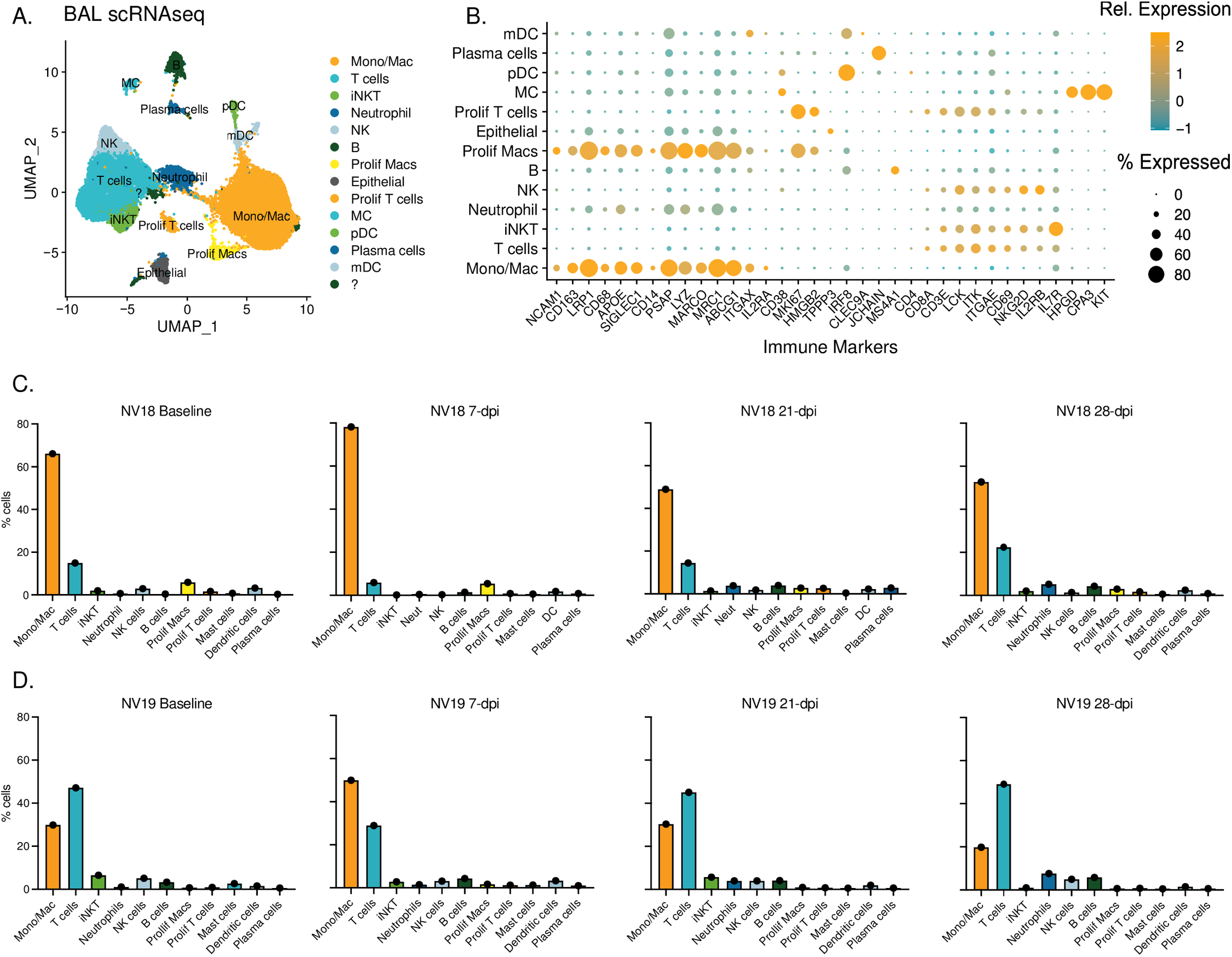
Single-cell classification and dynamics of bronchoalveolar lavage cell populations during SIV/SARS-CoV-2 coinfection. **A.** UMAP plots illustrating scRNAseq data obtained from BAL sampling of PTMs (NV18 and NV19) coinfected with SIV and SARS-CoV-2. **B.** Gene markers utilized for cell type identification. Dot color represents relative gene expression (Rel. Expression), while dot size indicates the proportion of cells expressing the gene (% Expression). Refer to supplemental figure 6 for additional genes used. **C-D.** Immune cell dynamics in BAL during SARS-CoV-2 infection for the more immunocompromised animal, NV18, (**C**) and NV19 (**D**). The baseline (BL) sample was collected prior to SARS-CoV-2 exposure at 48 weeks post-SIV infection. MC = Mast cells, pDC = plasmacytoid dendritic cells, mDC = myeloid dendritic cells, NK = natural killer cells

Prior to SARS-CoV-2 challenge, the more immunocompromised animal (NV18) exhibited a higher proportion of monocytes/macrophages in the lung, while the other coinfected animal (NV19) had higher levels of T cells (Figs 8C and 8D). At 7-dpi, both animals experienced an increase in monocytes/macrophages compared to baseline. Notably, the more immunocompromised animal consistently had higher levels of proliferating macrophages at all timepoints. We also observed an increase in neutrophils late in infection at 28-dpi. Following an initial decrease in the proportion of T cells at day 7, T cell levels rebounded to baseline by day 21, with a slight increase observed at 28-dpi. Additionally, B cell levels peaked in both animals at 28-dpi.

### Single-Cell Sequencing Identifies Multiple Cell Types Containing Viral RNA

We identified a diverse range of cell types containing viral transcripts by aligning the sequencing reads to the macaque, SARS-CoV-2, and SIV genomes (S6 and S7 Figs). Interestingly, we found SARS-CoV-2 RNA predominantly in DCs (S6B and S6C Figs), while neutrophils contained the highest percentage of SIV RNA (S7B and S7C Figs). The presence of vRNA in these cells can be attributed to various factors, including active viral replication, phagocytosis of infected cells, or potential viral contamination during the processing stages involved in single-cell sequencing [47]. We detected SARS-CoV-2 RNA exclusively at 2-dpi (S6D Fig), whereas SIV RNA was detectable at baseline and on days 21 and 28, indicating ongoing viral activity during those timepoints (S7D Fig). It is important to note that due to sample limitations, we only have scRNAseq data for one animal at 2-dpi. Therefore, we excluded the 2-dpi timepoint from any further analysis.

### scRNAseq Reveals Diverse Monocyte/Macrophage Populations in BAL

We then used single-cell analysis to gain a deeper understanding of monocyte/macrophage dynamics in BAL during SARS-CoV-2 infection in SIV+ PTMs. Specifically, we performed a subclustering analysis of the “Mono/Mac” cluster depicted in Figure 8A. This analysis revealed six subclusters characterized by variable gene expression patterns (Figs 9, S8A, and S8B). Among these, four populations exhibited features suggestive of alveolar macrophages, while the remaining two displayed characteristics associated with infiltrating/monocyte-derived macrophages. Typical markers of alveolar macrophages include CD68, CD11b (*ITGAM*), CD206 (*MRC1*), and the scavenger receptor class-A marker (*MARCO*). Within these subclusters, we identified resting macrophages [48] (*FABP4+ DDX60-*), infiltrating monocytes, anti-inflammatory macrophages [49] (*APOE*^hi^), *FPR3*^hi^ macrophages, activated macrophages (*IDO1^hi^, CXCL10^hi^*), and proliferating macrophages (*MKI67+, HMGB2*+). The more immunocompromised animal, NV18, exhibited a prominent increase in monocyte-derived cells with a more inert phenotype at 7-dpi, rising from 23% prior to SARS-CoV-2 infection to 60% of the total monocyte/macrophage population. This elevation persisted over the remaining 28 days of sampling (Fig 9C, S8A and S8B). While monocyte-derived cells dominated the pulmonary immune landscape of NV18, NV19 demonstrated an increase not only in monocyte-derived cells but also in anti-inflammatory macrophages (*APOE*^hi^), activated macrophages (*IDO1^hi^, CXCL10^hi^*), and proliferating macrophages at 7-dpi. Levels of all monocyte/macrophage subtypes began to normalize over time in this animal, with only anti-inflammatory macrophages remaining elevated at 28-dpi.

**Figure 9.**
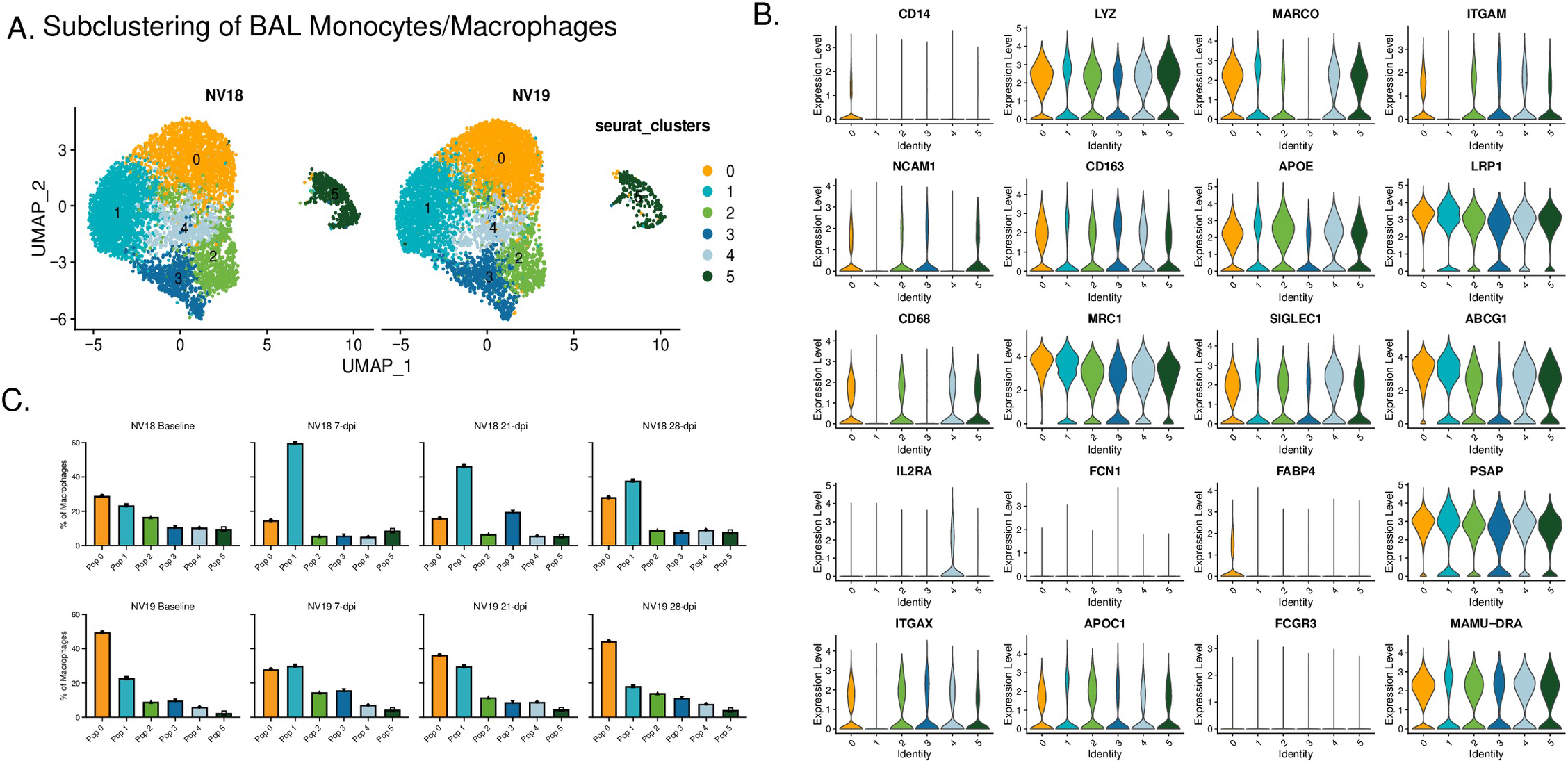
Monocyte/macrophage dynamics following SARS-CoV-2 inoculation of SIV+ PTM. **A.** UMAP plots of pulmonary monocyte/macrophage (mono/mac) subclustering. **B.** Violin Plots illustrating expression of canonical monocyte/macrophage gene signatures among the Seurat-derived clusters. **C.** Monocyte/macrophage dynamics during SARS-CoV-2 infection.

To gain additional insights into the monocyte/macrophages during coinfection, we performed gene set enrichment analysis (GSEA) on differentially expressed genes (DEGs) comparing the two coinfected animals at baseline and 7-dpi (Figs S9A and S9B). Prior to SARS-CoV-2 infection, NV18 exhibited enrichment in IFN-γ and IFN-⍺ responses, indicating greater activation of these pathways, potentially due to elevated SIV viremia and SIV-associated disease severity in this animal (S9A Fig). However, at 7-dpi, monocytes/macrophages in the other animal (NV19) showed enrichment in pathways typically upregulated during a respiratory infection, such as TNF-⍺ and IL-6 signaling, inflammatory response, complement, and coagulation (S9B Fig). Using GSEA to capture dynamic changes in monocyte/macrophage functionality, we incorporated Hallmark and KEGG terms and compared gene sets at baseline to gene sets at 7, 21, and 28-dpi for each animal (S9C Fig). Considering the substantial influx of monocytes with a less activated phenotype in the more immunocompromised animal (NV18) (Figs 9C and S8A-B), it was not surprising that GSEA comparing post-infection to baseline revealed decreased enrichment in the majority of pathways examined (S9C Fig). NV18 also exhibited a decrease in the frequency of CD169+ monocytes/macrophages on days 7 and 21 post-infection, further illustrating the limited functionality of the infiltrating monocytes in this animal. In contrast, NV19 followed a more typical pattern with DEGs enriched in the inflammatory response, cytokine and chemokine signaling, phagocytosis, and proliferation at 7-dpi. Though we found no evidence of actively replicating virus at the time, GSEA of days 21 and 28 post-infection revealed a continued enrichment of pathways associated with the inflammatory response in NV19.

### T cell Dynamics in Coinfected Animals

We also examined the T cell dynamics and phenotypes in the coinfected animals (Fig 10). Using subclustering analysis, we identified five distinct T cell subclusters, each with unique phenotypic characteristics. Populations 0, 2, and 3 had elevated expression of *CD69* and *ITGAE* (CD103) (Fig 10D), indicative of a tissue-resident phenotype (T_RM_). Cluster 0 displayed a more cytotoxic phenotype, characterized by elevated expression of *KLRD1, GZMB*, and *GZMK* (Fig 10C). Cluster 3 demonstrated an inflammatory phenotype, with greater expression of *IFN-γ* and tumor necrosis factor (TNF) cytokines *TNF* and *TNFSF8*, while cluster 2 represented an intermediate phenotype (Fig 10C). Additionally, we identified infiltrating T cells (cluster 1) and proliferating T cells (cluster 4, *MKI67^hi^* and *HMGB2^hi^*) (Fig 10D).

**Figure 10.**
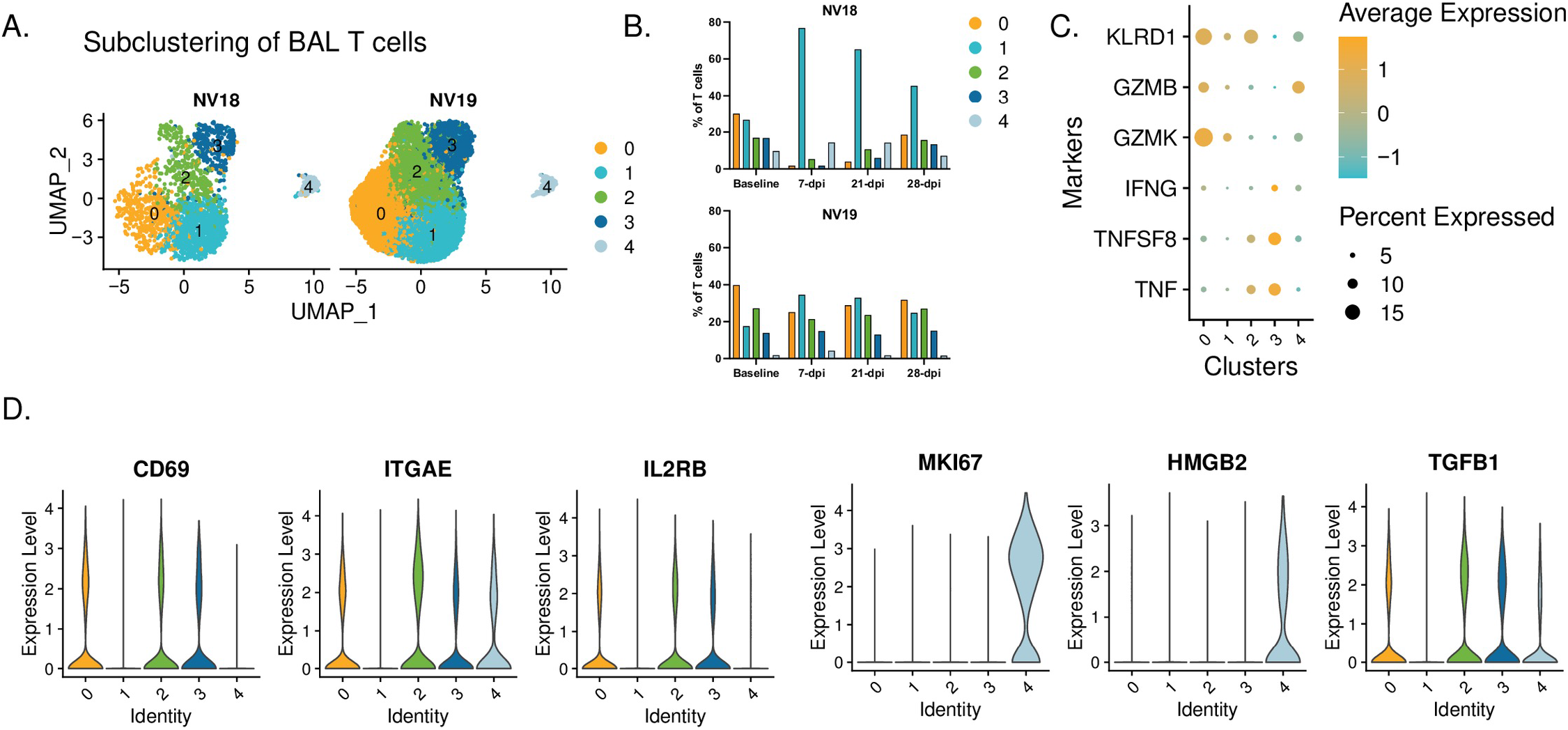
T cell dynamics. **A.** UMAP plots showing subclustered T cells. **B.** T cell dynamics during SARS-CoV-2 infection. **C.** Dot plot depicting proinflammatory cytokine and cytotoxic marker expression by cluster. **D.** Violin plots depicting gene expression of T cell markers in Seurat-derived T cell clusters.

Coinciding with the substantial influx of less activated monocytes at 7 dpi, the more immunocompromised animal, NV18, also experienced a notable shift in T cells towards a more inert phenotype (cluster 1). This population dominated the T cell landscape and persisted as the major population throughout the 28-day post-infection period (Fig 10B). Moreover, NV18 displayed increases in the proportion of proliferating T cells (cluster 4) at 7 and 21-dpi, indicating active cellular proliferation, accompanied by a substantial decrease in the proportions of all three T_RM_ clusters. Our flow cytometry results indicated that NV18 had a reduction in pulmonary CD3+ T cells following SARS-CoV-2 exposure. Thus, the T cell patterns observed in our scRNAseq analysis for this animal most likely reflect the preservation of specific subpopulations rather than actual increases. In contrast, at 7-dpi, NV19 displayed an increase in the proportion of infiltrating T cells (cluster 1) and a reciprocal decrease in T_RM_ cluster 0, although to a lesser extent than the more immunocompromised animal (NV18). Notably, all other T cell populations remained fairly stable in NV19.

## Discussion

Since its emergence in December 2019 in Wuhan, China, the novel coronavirus SARS-CoV-2 has had a profound global impact [1]. COVID-19, caused by SARS-CoV-2, encompasses a spectrum of disease manifestations, ranging from asymptomatic [50,51] to mild flu-like symptoms to pneumonia [52,53]. While the majority of infected individuals exhibit mild to moderate symptoms, a select group can experience severe complications marked by significantly elevated levels of coagulation biomarkers and proinflammatory cytokines, which can lead to acute respiratory distress syndrome (ARDS), and in some cases, death [54]. Risk factors such as a compromised immune system, advanced age, and comorbidities such as cardiovascular disease, diabetes, and obesity increase the likelihood of severe disease.

The presence of HIV infection poses additional risks for individuals, including a compromised immune system and a higher prevalence of cardiovascular disease, raising concerns about the impact of HIV on the severity and persistence of SARS-CoV-2 infections [13–15]. While initial research indicated similar or improved outcomes for people living with HIV (PLWH) compared to the general population [16–18], larger population-based studies reported higher rates of hospitalization and COVID-19-related deaths among PLWH [19–23]. Recent studies suggest that unsuppressed viral loads or low CD4+ T cell counts are associated with suboptimal cellular and humoral immune responses to SARS-CoV-2 [24,25].

Immunodeficiency associated with HIV not only raises concerns about increased severity but also the potential facilitation of SARS-CoV-2 persistence and evolution, leading to the emergence of novel viral variants. Karim et al, (2021) highlighted this concern in a recent study in which an individual with advanced HIV showed prolonged shedding of SARS-CoV-2, high viral loads, and the development of multiple viral mutations [26]. Although retrospective studies have explored the impact of HIV status on COVID-19 incidence and severity, controlled studies in this area are lacking.

To address these gaps, we conducted a small pilot study involving two pigtail macaques (PTMs) infected with SIVmac239, a strain that is highly pathogenic in PTM and models progressive HIV infection, and subsequently exposed them to SARS-CoV-2 after approximately one year. Notably, PTMs infected with SIV exhibit more rapid progression to AIDS as compared to rhesus macaques and demonstrate cardiovascular abnormalities similar to those observed in humans with advanced HIV, making them an ideal model for evaluating the effects of chronic SIV infection on SARS-CoV-2 dynamics [55–58]. Our study aimed to investigate the impact of SIV-induced immunodeficiency on the clinical manifestation of COVID-19, along with its impacts on viral replication and evolution in a controlled setting. We compared the clinical, virological, and immunological outcomes of the coinfected animals with our previously published cohort of SIV-naïve PTMs infected with SARS-CoV-2 [27].

One of the key findings of our study is that SIV-induced immunodeficiency did not lead to enhanced COVID-19 disease in the coinfected animals. Despite the presence of significant immunodeficiency, as evidenced by the severe reduction in CD4+ T cells, the coinfected animals exhibited only mild COVID-19 symptoms, similar to the control group. This finding contrasts with previous studies that have reported a higher risk of severe disease and mortality in PLWH [19–23], suggesting that aspects beyond immunodeficiency, such as comorbidities or host-related factors, may contribute to the elevated risk of severe COVID-19 observed in PLWH.

Our analysis of SARS-CoV-2 viral dynamics in the coinfected animals revealed that SIV-induced immunodeficiency did not significantly impact viral replication or evolution, with viral dynamics indistinguishable from the controls. Despite higher levels of vRNA in bronchoalveolar lavage (BAL) of the more immunocompromised animal (NV18), vRNA levels became undetectable in both of the coinfected animals by three- or four-weeks post-infection in all sampled mucosal sites indicating that underlying SIV infection alone is insufficient to drive uncontrolled SARS-CoV-2 replication.

However, we did observe a notable difference in the adaptive immune response to SARS-CoV-2 infection between the SIV+ and SIV-naïve PTMs. By 21-dpi, the control animals exhibited detectable SARS-CoV-2-specific neutralizing antibodies, IgA and IgG binding antibodies, and virus-specific T cell responses. In contrast, both coinfected animals failed to generate virus-specific humoral or cellular immune responses against SARS-CoV-2. This finding is consistent with studies linking uncontrolled HIV infection to suboptimal T cell and antibody responses to SARS-CoV-2 [24,25]. These results underscore the impact of pre-existing immunodeficiency on the development of adaptive immunity during coinfection. The observed inability to mount effective virus-specific cellular and humoral immune responses sheds light on the potential challenges faced by individuals with advanced HIV infection when encountering SARS-CoV-2 and raises concerns about the potential impacts of reinfection.

Additionally, we noted an influx of pulmonary infiltrating monocytes, which dominated the immunological landscape in the coinfected animals, compared to T cells in SIV-naïve animals, as reported in our previous study. Our scRNAseq analysis provided additional insights into nuances of the immune response in the coinfected PTMs, particularly in the more immunocompromised animal (NV18), which exhibited an influx of pulmonary monocytes and T cells with diminished functionality. Immune dynamics in the less immunocompromised animal (NV19) suggested a more balanced immune response.

The observed differences in pulmonary infiltrating immune cells between the two coinfected animals may be attributed to the varying levels of viremia, overall immune competence, or subclinical pneumocystis infection in the more immunocompromised animal (NV18).

## Conclusion

Overall, our study provides valuable insights into the interplay between SIV-induced immunodeficiency and SARS-CoV-2 infection. Despite the notable immunodeficiency observed in the coinfected animals, we found no evidence of enhanced COVID-19 disease nor significant impacts on viral replication or evolution. However, the impaired T-cell response and lack of neutralizing antibodies in the coinfected animals highlight the impact of underlying SIV-induced immunodeficiency on the immune response to SARS-CoV-2. These findings contribute to our understanding of COVID-19 pathogenesis in immunocompromised individuals and may help guide the development of strategies to manage COVID-19 in vulnerable populations.

## Limitations

As this was a preliminary study involving only two animals, it will be necessary to conduct follow-up studies with larger cohorts in order to validate our findings. Nonetheless, our data provide novel important discoveries contributing to the growing collection of SARS-CoV-2 resources. Further investigations into SARS-CoV-2 reinfection of SIV+ nonhuman primates could serve as a promising follow-up to this study. Our coinfection model demonstrated that the innate immune response was likely efficient in eliminating SARS-CoV-2 infection. A study that compares reinfection rates and viral clearance upon secondary exposure would be an exciting next avenue to pursue.

## Acknowledgements

Work described in this report was conducted at the Tulane National Primate Research Center (RRID: SCR_008167) with support from TNPRC core labs (RRIDs: SCR_024606, SCR_024609, SCR_024613, SCR_024611, SCR_024612, SCR_024614, SCR_024610, SCR_024679). Pigtail macaques were purpose bred at the Johns Hopkins University breeding colony supported by U42OD013111. Work was supported by instrumentation support funds NIH S10 OD026800 and NIH S10 OD030347. The following reagent was obtained from BEI Resources; SARS-CoV-2 isolate WA1/2020 (NR-52281). We thank Kejing Song for help with sequencing, and Ann-Marie May and Carli Thompson from the Tulane Office of Biosafety for oversight of experiments in BSL-3 containment.

---

**Supplemental Figure 1.**
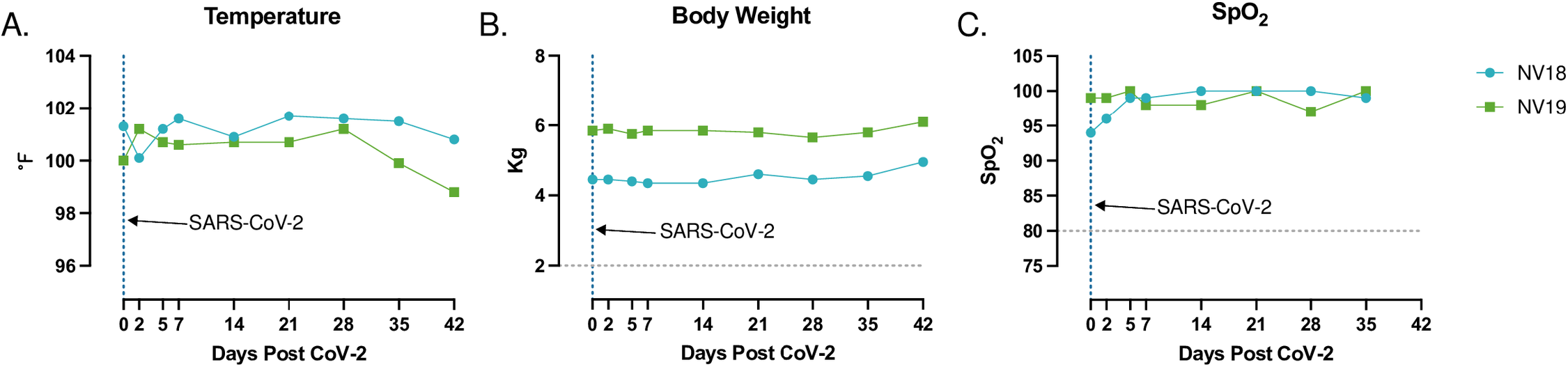
Temperature (**A**), weight (**B**), and saturation of peripheral oxygen (SpO2) (**C**) levels were measured prior to and for 6 weeks following SARS-CoV-2 inoculation of SIV+ pigtail macaques. Day 0 indicates time of SARS-CoV-2 infection, 48 weeks post SIVmac239 exposure.

**Supplemental Figure 2.**
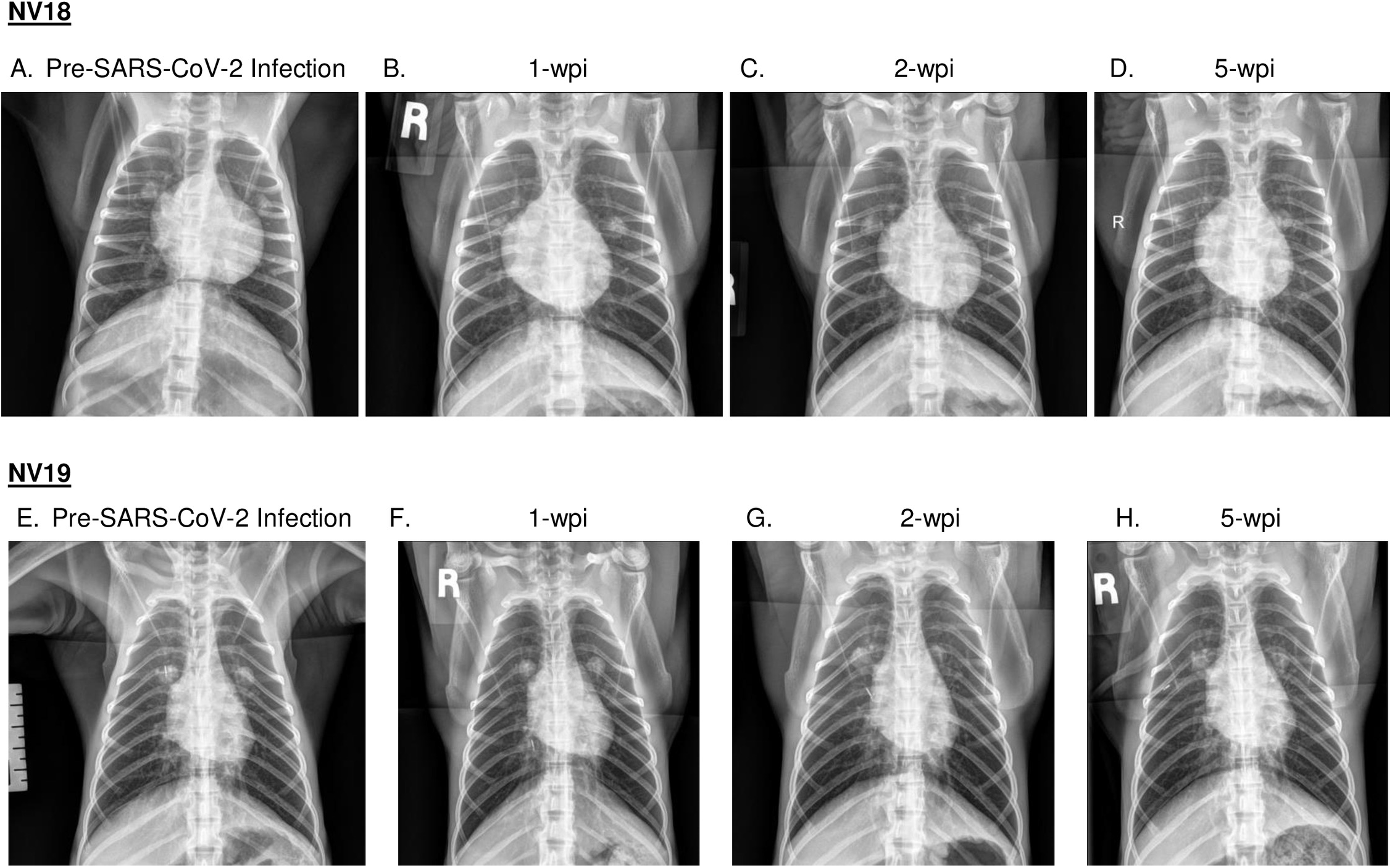
Radiographs of SIV-infected pigtail macaques (PTM) challenged with SARS-CoV-2. Radiographs were obtained prior to SARS-CoV-2 infection and at weeks 1-, 2-, and 5-weeks post infection (wpi). Baseline was established at 2 weeks prior to SARS-CoV-2 inoculation.

**Supplemental Figure 3.**
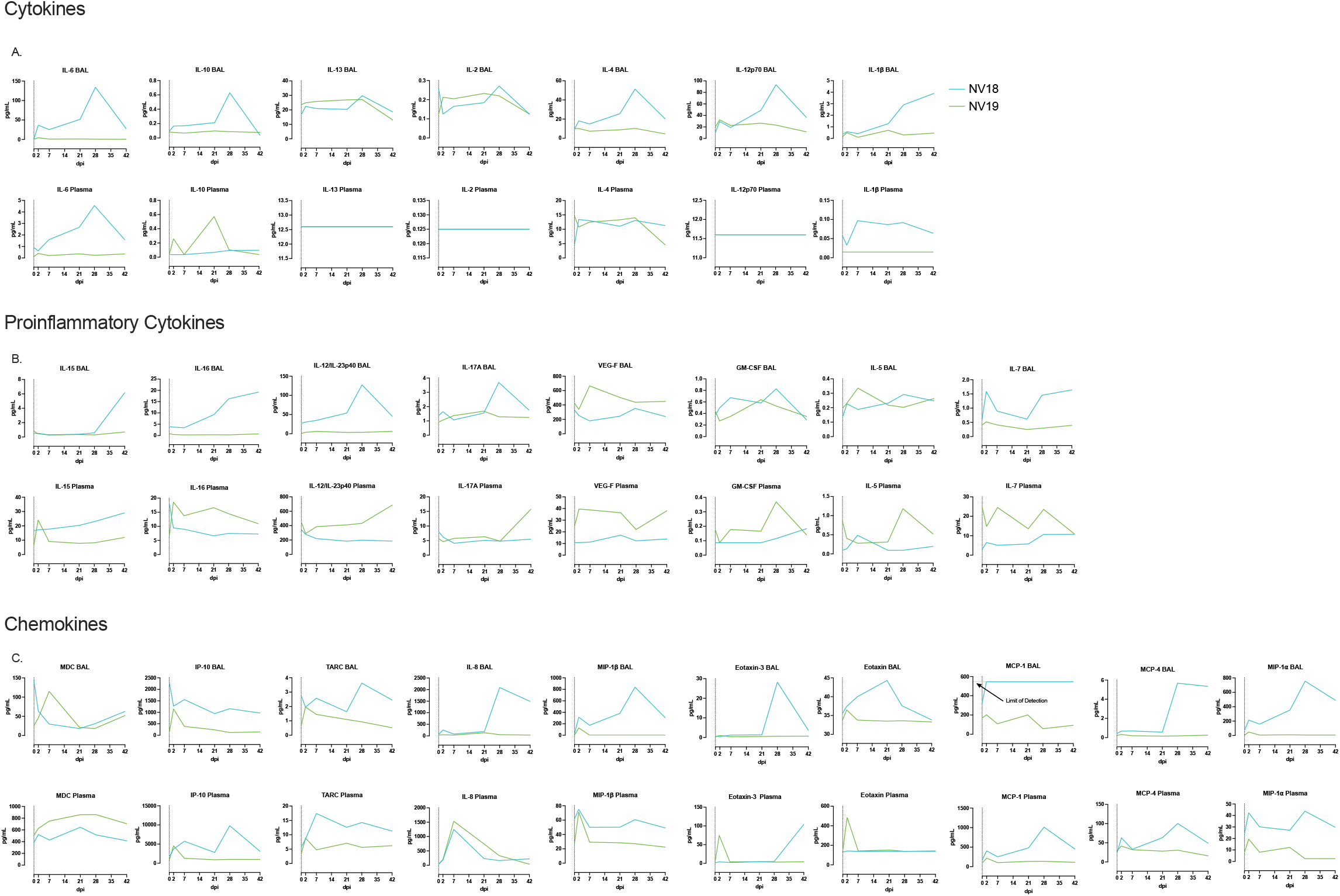
Meso Scale analysis of cytokine and chemokine fluctuations in blood and BAL in PTM coinfected with SIV and SARS-CoV-2. **A-C.** Line graphs illustrating cytokine, proinflammatory cytokine, and chemokine dynamics in BAL supernatant and plasma before and 2-, 7-, 21-, 28-, and 42-days post SARS-CoV-2 infection.

**Supplementary Figure 4.**
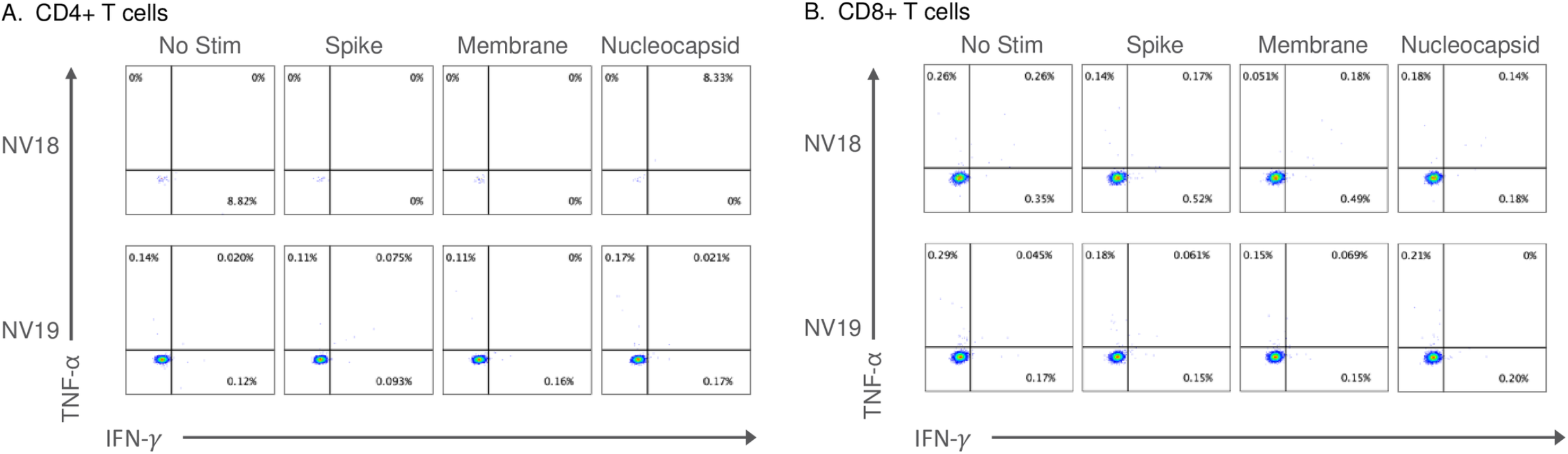
Peripheral SARS-CoV-2 specific T cell responses were undetectable 21-days-post-infection. Two female pigtail macaques (PTM, NV18 & NV19) co-infected with SIVmac239 and SARS-CoV-2 shown. **A&B**. Flow cytometry dot plots demonstrating the IFN-γ and TNF-⍺ response of CD4+ (**A**) and CD8+ (**B**) T cells to overnight SARS-CoV-2 peptide (spike, membrane, and nucleocapsid) stimulation. No Stim = cells incubated overnight without peptide stimulation.

**Supplementary Figure 5.**
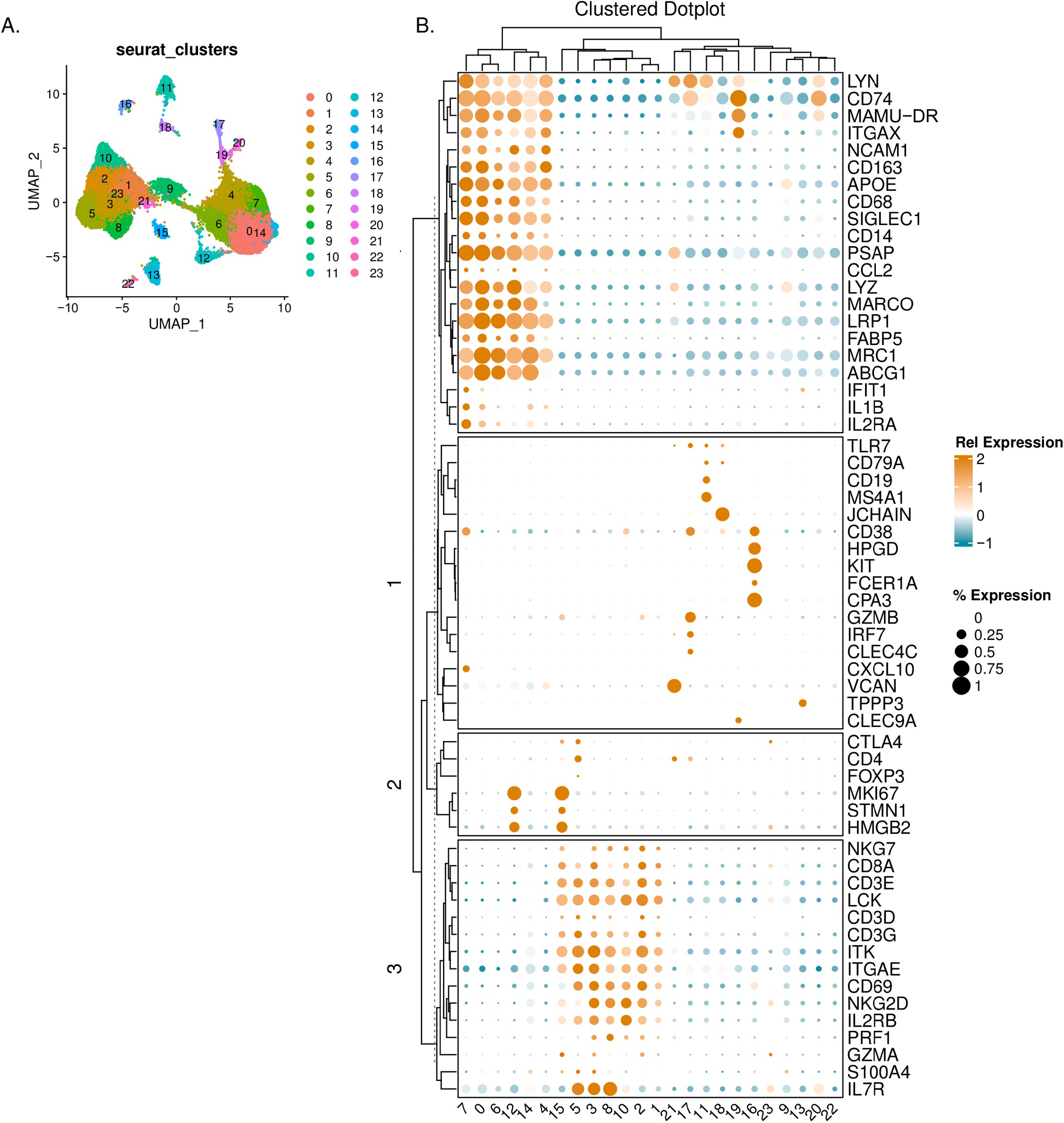
Defining Seurat clusters. **A.** UMAP displaying Seurat-derived clusters obtained from FindAllMarkers function in Seurat. **B.** Dot plot depicting markers used to identify cell types. Hierarchical clustering method (hclust) was used to cluster columns and rows. The color of the dots indicates the relative gene (Rel. Expression). Dot size represents the percentage of cells expressing the gene (% Expression).

**Supplemental Figure 6.**
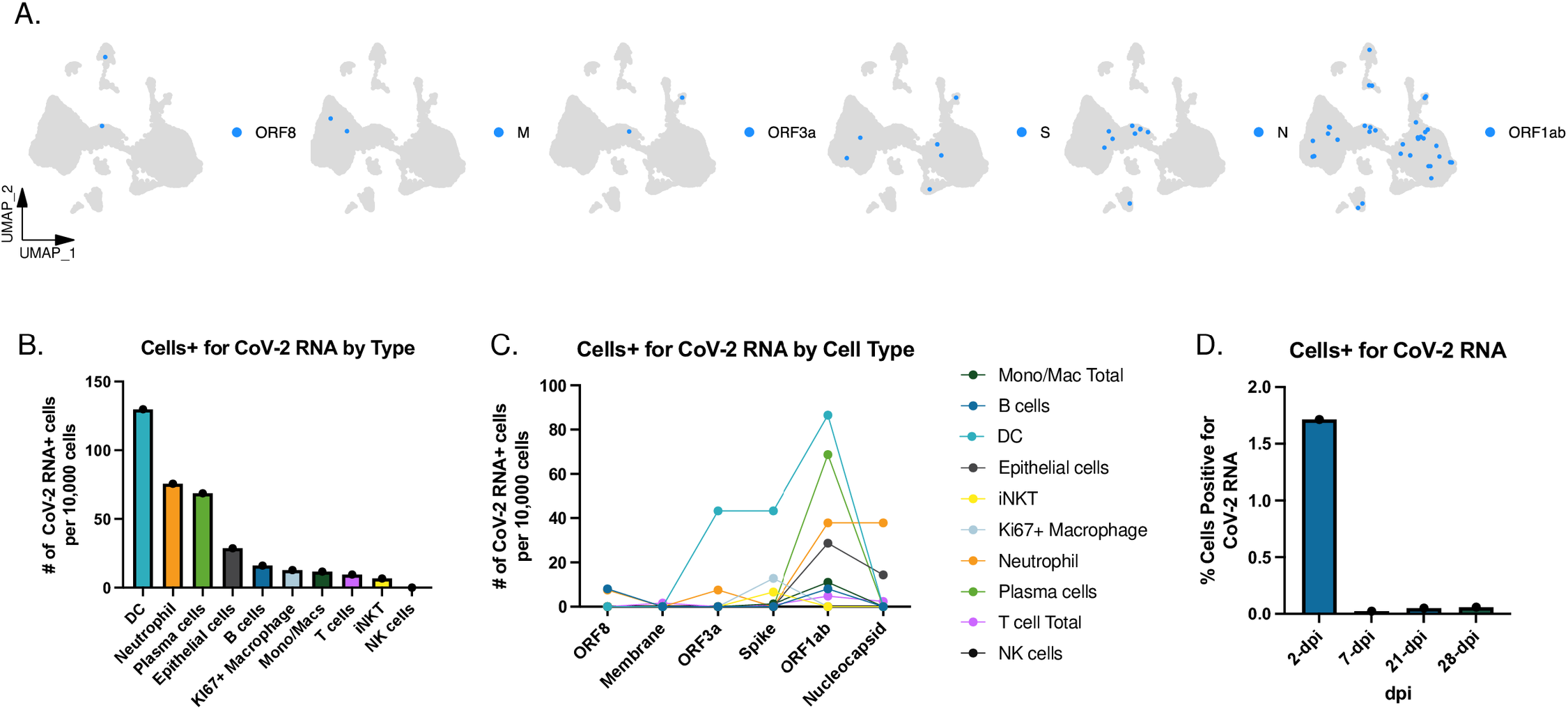
Single-cell analysis of SARS-CoV-2 positive cells in BAL of SIV+ PTM. **A.** UMAP plots highlighting cells with detectable SARS-CoV-2 transcripts. **B.** Distribution of cells by cell type positive for any SARS-CoV-2 transcript. **C.** Number of cells positive for specific SARS-CoV-2 transcripts grouped by cell type. **D.** Percentage of cells positive for SARS-CoV-2 transcripts by days post-infection (dpi).

**Supplemental Figure 7.**
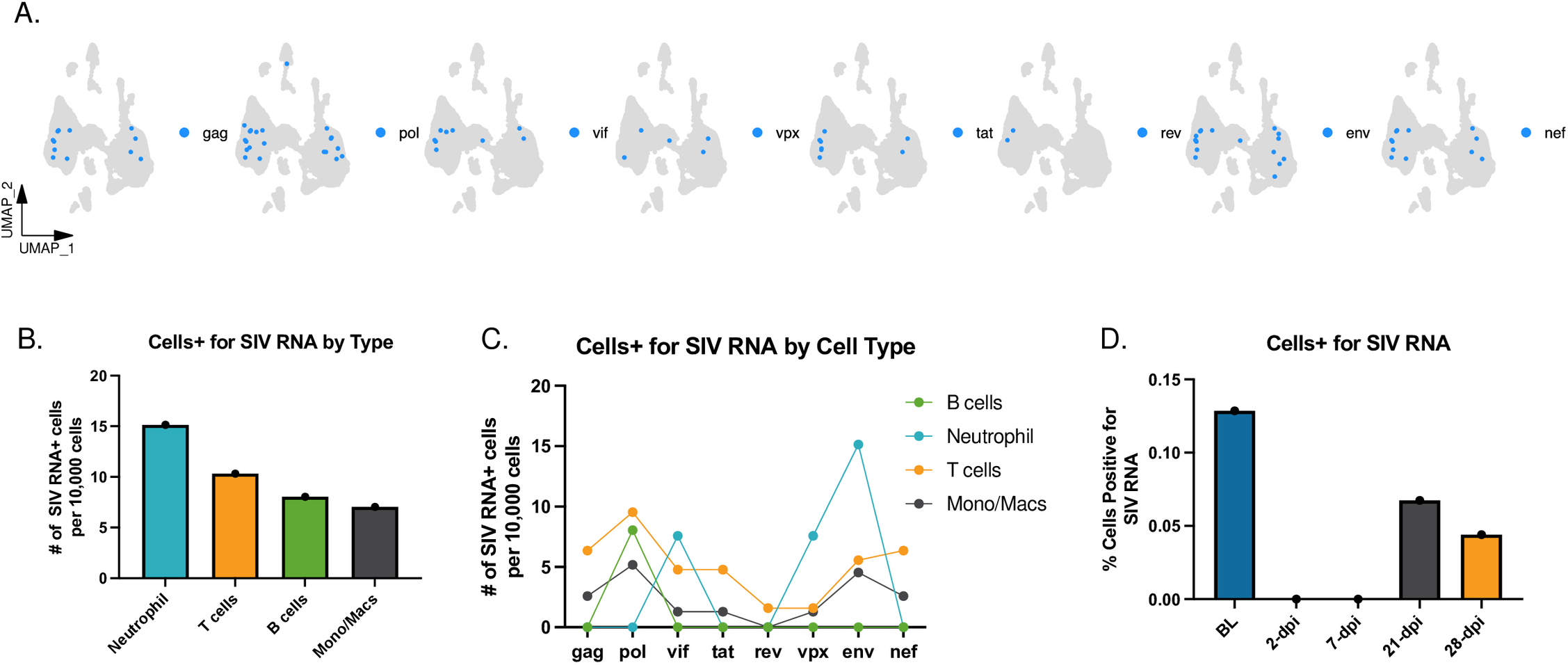
Single-cell analysis of SIV positive cells in BAL of coinfected PTM. **A.** UMAP plots highlighting cells with detectable SIV transcripts. **B.** Distribution of cells grouped by type exhibiting any SIV transcripts. **C.** Number of cells positive for specific SIV transcripts grouped by cell type. **D**. Percentage of cells positive for SIV transcripts by days exposure.

**Supplementary Figure 8.**
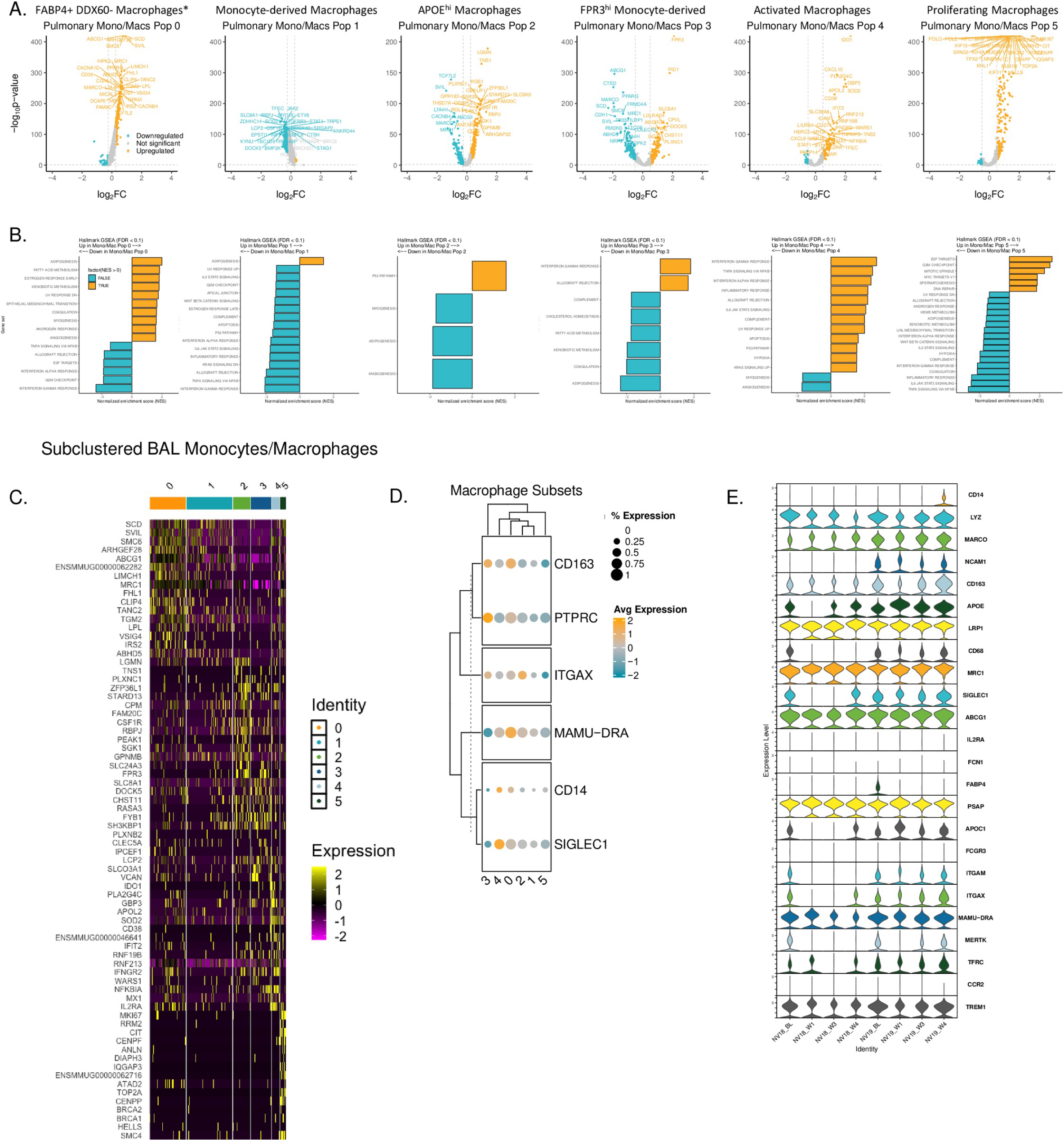
Single-cell monocyte/macrophage characterization. **A.** Volcano plots displaying significantly upregulated and downregulated differentially expressed genes (DEGs) in the Monocyte/macrophage populations. **B.** Bar plots depicting normalized net enrichment score (NES) from gene set enrichment analysis (GSEA) of Hallmark biological pathways. A false discovery rate (FDR) cutoff of 0.1 was used to determine significance. **C.** Heatmap of top 10 differentially expressed genes for each monocyte/macrophage cluster. **D.** Dot plot illustrating gene expression differences among the Seurat-derived clusters, corresponding to markers used in flow cytometry analysis (Figure 4 panel K). Hierarchical clustering (hclust) was used for column and row clustering. Dot color represents relative gene expression (Rel Expression), while dot size indicates the percentage of cells expressing the gene (% Expression). **E.** Stacked violin plot illustrating gene expression patterns of monocytes/macrophages at baseline (BL) and weeks 1, 3, and 4 post-SARS-CoV-2 infection.

**Supplemental Figure 9.**
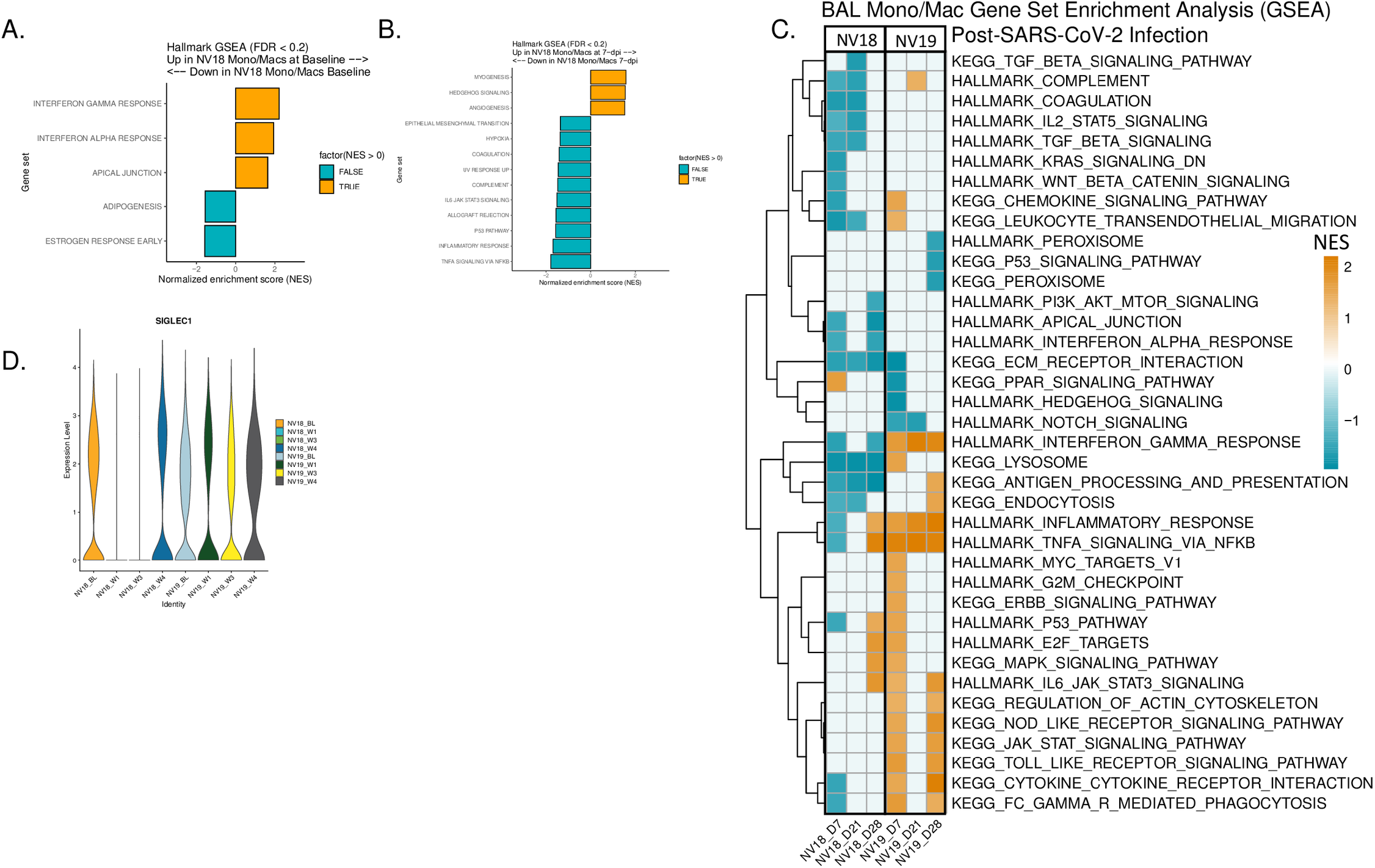
Differential gene expression (DEG) and enrichment analysis of Monocyte/Macrophage Subsets in NV18 and NV19. **A&B.** Gene set enrichment analysis (GSEA) of DEGs comparing NV18 and NV19 at baseline (**A**) and 7-dpi (**B**). **C.** GSEA results comparing DEGs at days 7, 21, and 28 post-exposure to baseline. A false discovery rate (FDR) cutoff of 0.2 was used to determine significance. **D.** CD169 (*SIGLEC1*) expression kinetics.

